# Trophic interaction modifications disrupt the structure and stability of food webs

**DOI:** 10.1101/345280

**Authors:** J. Christopher D. Terry, Rebecca J. Morris, Michael B. Bonsall

## Abstract

Trophic interaction modifications, where a consumer-resource interaction is influenced by an additional species, are established as being prevalent throughout ecological networks. Despite this, their influence on the structure of interaction distributions within communities has not yet been examined. Although empirical information about the distribution of interaction modifications is currently sparse, the non-trophic effects they induce will be structured by the underlying network of trophic interactions. Here we test the impact of interaction modifications, introduced under a range of distributional assumptions to artificial and empirical trophic networks, on the overall structure of interactions within communities. We show that local stability and reactivity is critically dependent on the inter-relationship between the trophic and non-trophic effects. Depending on their distribution, interaction modifications can generate significant additional structure to community interactions making analyses of the stability of ecological systems based solely on trophic networks unreliable. Empirical information on the topological and strength distributions of interaction modifications will be a key part of understanding the dynamics of communities.

## Introduction

Understanding how large and diverse ecosystems per
sist is a core challenge in ecology. Theoretical expectations that large random complex systems are unlikely to be locally stable (May 1972), indicate that the interactions within ecological communities are structured in important ways (Montoya *et al.* 2006). While network analyses need not be inherently specific to a particular class of interaction (May 1972, Allesina and Tang 2012), trophic interactions are commonly focussed upon because of their role in determining energy flux and the comparatively high level of empirical understanding. A growing body of work has demonstrated features of trophic networks that can stabilise communities - including the distribution of weak links (McCann *et al.* 1998, Neutel *et al.* 2002), pairwise correlations (Tang *et al.* 2014), modularity (Grilli *et al.* 2016), row structure (Jacquet *et al.* 2016) and trophic level coherence (Johnson and Jones 2017). However, ecological communities contain complex networks of interactions beyond trophic relationships (Ings *et al.* 2009), and there is an emerging appreciation of the value in studying the full spectrum of interaction types simultaneously (Olff *et al.* 2009, Fontaine *et al.* 2011, Kéfi *et al.* 2012, Mougi and Kondoh 2012, Coyte *et al.* 2015, Pilosof *et al.* 2017, García-Callejas *et al.* 2018).

The ubiquity and diversity of non-trophic interactions poses a considerable problem for ecologists as organising features appear few and far between. However, networks of non-trophic effects do show considerable non-random structure (Kéfi *et al.* 2016). A notable fraction of non-trophic effects are caused by interaction modifications (Kéfi *et al.* 2012), where a pairwise interaction is dependent on a third species (Wootton 1994). These are pervasive within ecological networks (Werner and Peacor 2003) and capable of exerting impacts as strong as direct trophic interactions (Preisser *et al.* 2005).

Classic examples of interaction modifications include shifts in foraging patterns in response to the threat of predators (Suraci *et al.* 2016) and species providing associational resistance to predation (Barbosa *et al.* 2009), but the concept also encompasses any biotic influences on foraging patterns such as the availability of alternative prey (Abrams 2010) and many ecosystem engineering effects (Sanders *et al.* 2014). This additional source of interactions and dynamic connectance introduces emergent relationships between species that may not otherwise directly interact, potentially greatly increasing the dynamic connectance of the system (Yodzis 2000). The incorporation of interaction modification effects has been empirically demonstrated to be necessary to understand species fitness (Mayfield and Stouffer 2017), how communities persist (van Veen *et al.* 2005), and the response of communities to species loss (Donohue *et al.* 2017).

Although the distribution of interaction modifications at the network level is essentially unknown, it is reasonable to expect that they will be structured, both internally and with respect to the underlying interactions being modified (Golubski *et al.* 2016). This relationship offers an opportunity to build upon the understanding of the structure of trophic networks (Dunne and Pascual 2006) to inform the likely distribution and consequences of non-trophic effects caused by trophic interaction modifications. While both trophic and non-trophic interactions can be subject to modifications (Kéfi *et al.* 2012), here we focus on trophic interaction modifications to start to move beyond random distributions of non-trophic effects (Mougi and Kondoh 2012, Coyte *et al.* 2015) and higher-order interactions (Wilson 1992, Bairey *et al.* 2016).

Here, we examine the impact non-trophic effects induced by trophic interaction modifications can have on the equilibrium dynamics of simple models of the interactions between species in artificial and empirical networks. We combine interaction matrices representing trophic networks with non-trophic effects generated with a variety of distinct interaction modification distribution models. We examine the properties of the resultant interaction matrices and test the impact on local stability and reactivity to show that the impact of interaction modifications depends crucially on their distribution and relationship with the underlying trophic network.

## Methods

Interaction modifications (Wootton 1994) are higher-order processes that induce non-trophic effects from the modifier species onto a pair of interactors. In this paper we distinguish between the underlying trophic interaction modification (TIM) process and the resultant (pairwise) non-trophic trophic effects (NTEs), also termed trait-mediated indirect interactions (Ohgushi *et al.* 2012). This distinction (Figure 1) between process and consequence aids the analysis of these complex processes (Terry *et al.* 2017).

**Figure 1:**
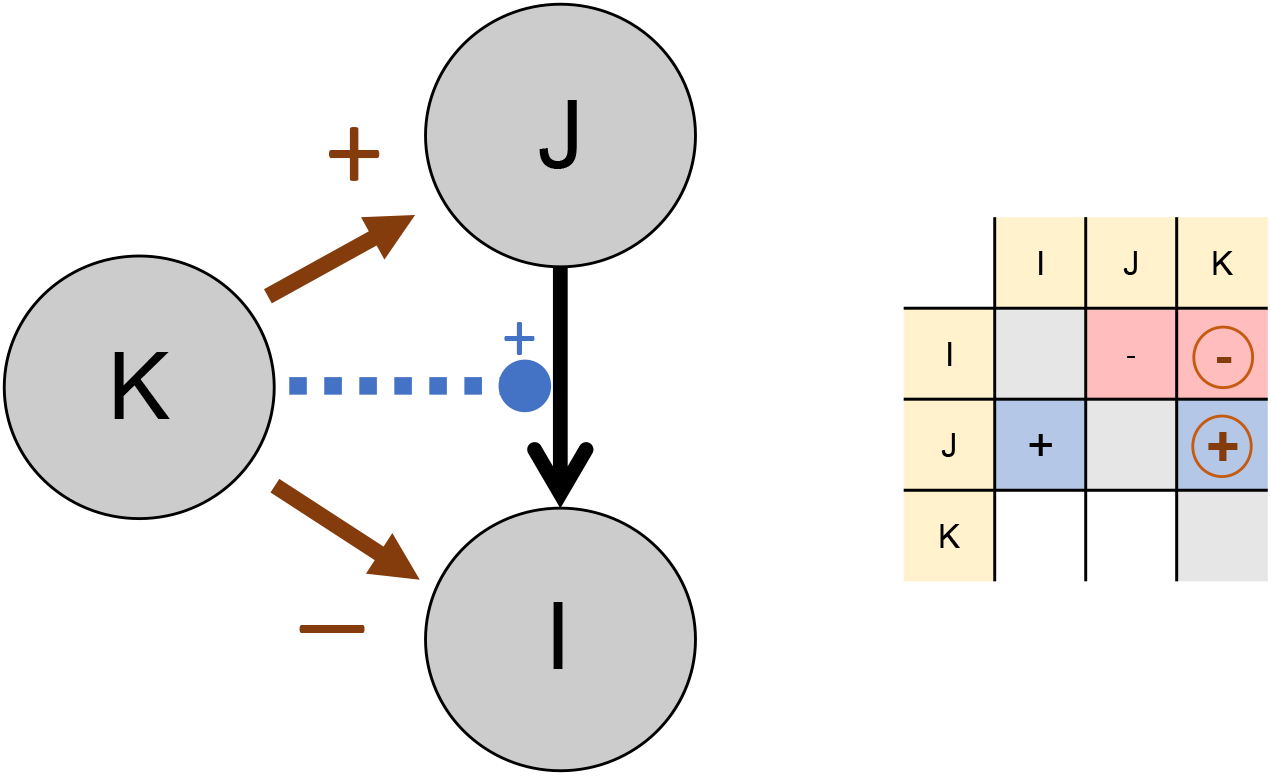
Schematic representation of the relationship between a trophic interaction modification (TIM, blue dashed line), consequent non-trophic effects (NTE, brown solid lines) and resultant community matrix. Each column of the community matrix shows the effect of an increase in a species on the growth rate of species in each row 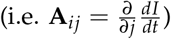. Here, an interaction modification results in two non-trophic effects (circled). The modifier species (*K*) acts to strengthen a trophic interaction between consumer *J* on resource *I.* This results in a positive NTE on *J* and a negative NTE on *I*.

### Communities as interaction matrices

We represent the complete set of interactions in a system as a Jacobian matrix **A**, specifically a community matrix (Novak *et al.* 2016), which is assumed to be derived from a set of populations each at a feasible equilibrium. Each element **A**_*ij*_, represents the instantaneous effect of a change in the population of species *j* on the population growth rate of species *i*. The community matrix is considered to be based on a linearisation of more complex processes that govern the relationships between the species and as such the description of the interactions is only strictly applicable close to the original non-trivial equilibrium. We determine **A** to be constructed from the combination of two matrices specifying the trophic (**B**) and non-trophic (**C**) interactions present in the community (Figure 2). For all matrices, we only consider inter-specific interactions, all intra-specific diagonal terms (**A**_*ii*_) were set to zero (see below).

**Figure 2:**
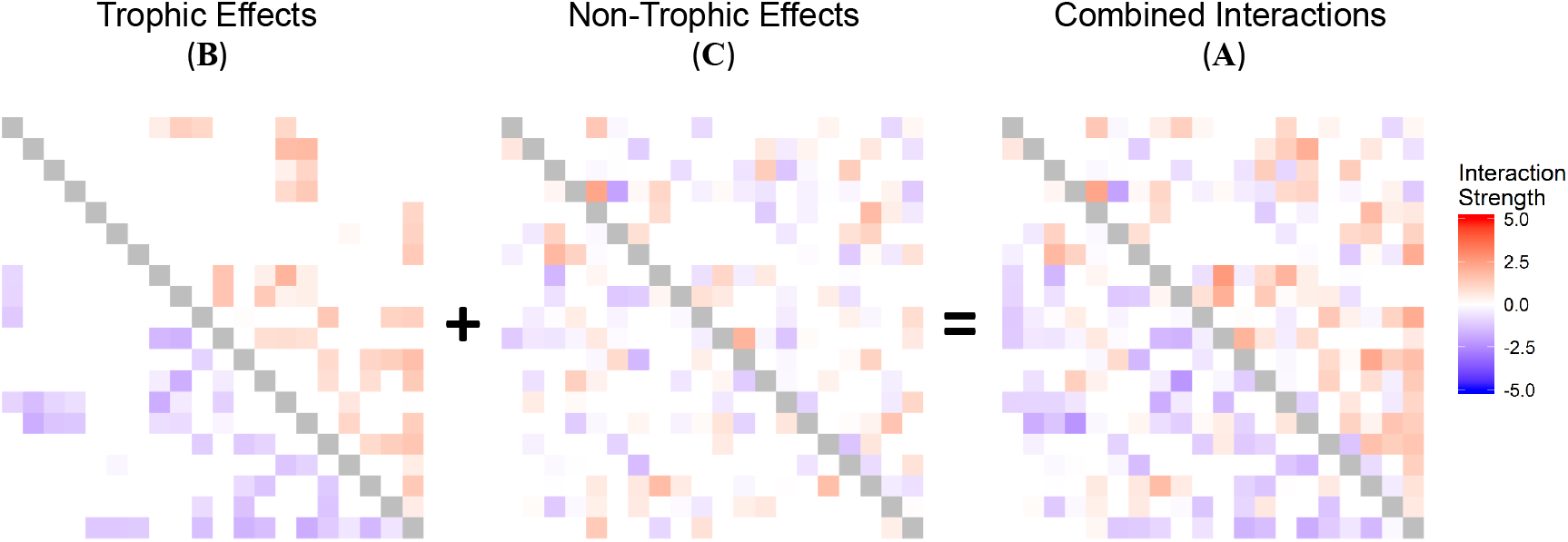
Illustration of construction of community matrices. At an assumed equilibrium, a combined community matrix (**A**) can be split into the impact of trophic (**B**) and non-trophic effects (**C**). The species are arranged approximately in trophic height order, with basal species top/left and top predators at the bottom/right. The underlying trophic network depicted was generated with the niche model (S=20, C=0.2), parameterised with a bivariate normal distribution *𝒩*(*μ_x_* = −1, *μ_y_* = 1, *sd_x_* = 0.5, *sd_y_* = 0.5, *ρ* = −0.8). Intra-specific interactions were fixed at zero and shown in grey.

### Generating trophic networks

We determined the trophic topology of the artificial communities using the niche model (Williams and Martinez 2000), a network generating algorithm that has been shown to reproduce many of the features of real food webs (Williams and Martinez 2008). We generated 100 networks each with 60 species and target connectance of 0.2. We parameterised these interactions with draws from a bivariate normal distribution *𝒩* (*μ_x_* = −1, *μ_y_* = 1, *sd_x_* = 0.5, *sd_y_* = 0.5, *ρ* = −0.8) to create each trophic interaction matrix **B**, following Allesina *et al.* (2015). This specification maintains a mean interaction value of 0 and an overall symmetry in impacts between consumer and resource. This simplifies analysis but is known to be unrealistic (Wootton and Emmerson 2005). We therefore repeated the analysis with increased consumer effects on resources (*μ_x_* = −5 & −10, SI 3). Since the draws are unbounded a small fraction of interactions had the ‘wrong’ sign combination for an exploitative interaction. As exploitative interactions still made up the overwhelming bulk of interactions, we did not remove these and for convenience we will refer to these underlying networks as trophic networks.

### Incorporating interaction modifications

Trophic interaction modifications are ‘higher-order interactions’ that act through at least three species (Terry *et al.* 2017). However, the short-term consequences of the interaction modification can be linearised to identify the effect of the modifier on the consumer and the resource (Wilson 1992, Bairey *et al.* 2016) at the system state under consideration (Figure 1). These non-trophic effects (NTEs) can be used to construct a matrix, **C**, of pairwise effects caused by the TIMs. In our representation the value of the original pairwise trophic interaction is left unchanged, i.e. it is assumed that the trophic interaction strengths in **B** already incorporate the consequence of the equilibrium level of the modifier species.

Based upon the trophic networks, NTE matrices were generated using a set of 16 models. These varied in topology, strength distribution and translation from TIM to NTEs, discussed below. As a baseline TIM model, each possible interaction modification (combination of consumer-resource pair and third modifier species) was equally likely to be present. Each TIM was assigned an effect parameter, *c_ijk_*, representing the size and direction of the modification by species *k* of the consumption of resource *i* by consumer *j*. These were specified such that the mean magnitude of individual NTEs (*α*) incorporated into a community is 0.5, by setting the standard deviation of the normal distribution to 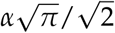 in line with results from meta-analysis that suggest an approximate correspondence between the strength of trophic and non-trophic interactions (Preisser *et al.* 2005). A positive *c_ijk_* indicates a facilitating modification where an increase in the modifying species would increase the strength of the interaction. It follows that this would lead to a positive effect of the modifier on the consumer and a negative effect of the modifier on the resource. A negative *c_ijk_* would cause the reverse, an interfering modification. The *c_ijk_* values determined the NTEs of the modifier on the two interactors (**C**_*jk*_= *c_ijk_*, **C**_*ik*_= −*c_ijk_*) and were used to construct a TIM effect matrix **C**.

We only considered TIMs where species modified the interaction between two other species - we did not allow species to modify their own interactions. Multiple NTE from one species to another were combined additively. TIMs were introduced at a TIM density (defined as the expected number of TIMs in the network per species, *ω*) of either 0, 1, 5, 10, 20, 30, 40 and 50. For a given trophic network, we identified each potential TIM that could exist following a particular distribution model, and assigned each potential TIM a probability of existing equal to 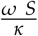, where *κ* is the total number of potential TIMs (i.e. combinations of consumer-resource and third species) and S the number of species. Note that TIM density as defined here is distinct to both ‘TIM connectance’ (the fraction of possible TIMs that are observed, which is dependent on the trophic connectance) and to non-trophic connectance (the resultant fraction of non-zero elements of the consequent TIM effect matrix **C**, which is dependent on the distribution and overlap of the TIMs).

### Different TIM models

Fourteen variations of the baseline TIM distributional model were examined, illustrated in Figure 3 with an example NTE matrix and a cartoon of the model’s distinguishing features. In addition, we also tested an NTE distribution model where there was no underlying interaction modification structure. There, NTEs were independently randomly distributed with their strengths drawn from a normal distribution. The mean magnitude of individual NTEs was kept constant across all models. In the results and discussion the key distinguishing feature is used to identify each model rather than a letter for clarity. The properties of empirical distributions of interaction modifications are unknown, and likely to significantly differ between communities. This set of models is therefore an attempt to examine the range of properties that such networks may have in order to identify how interaction modifications may introduce additional significant structure to ecological networks.

**Figure 3:**
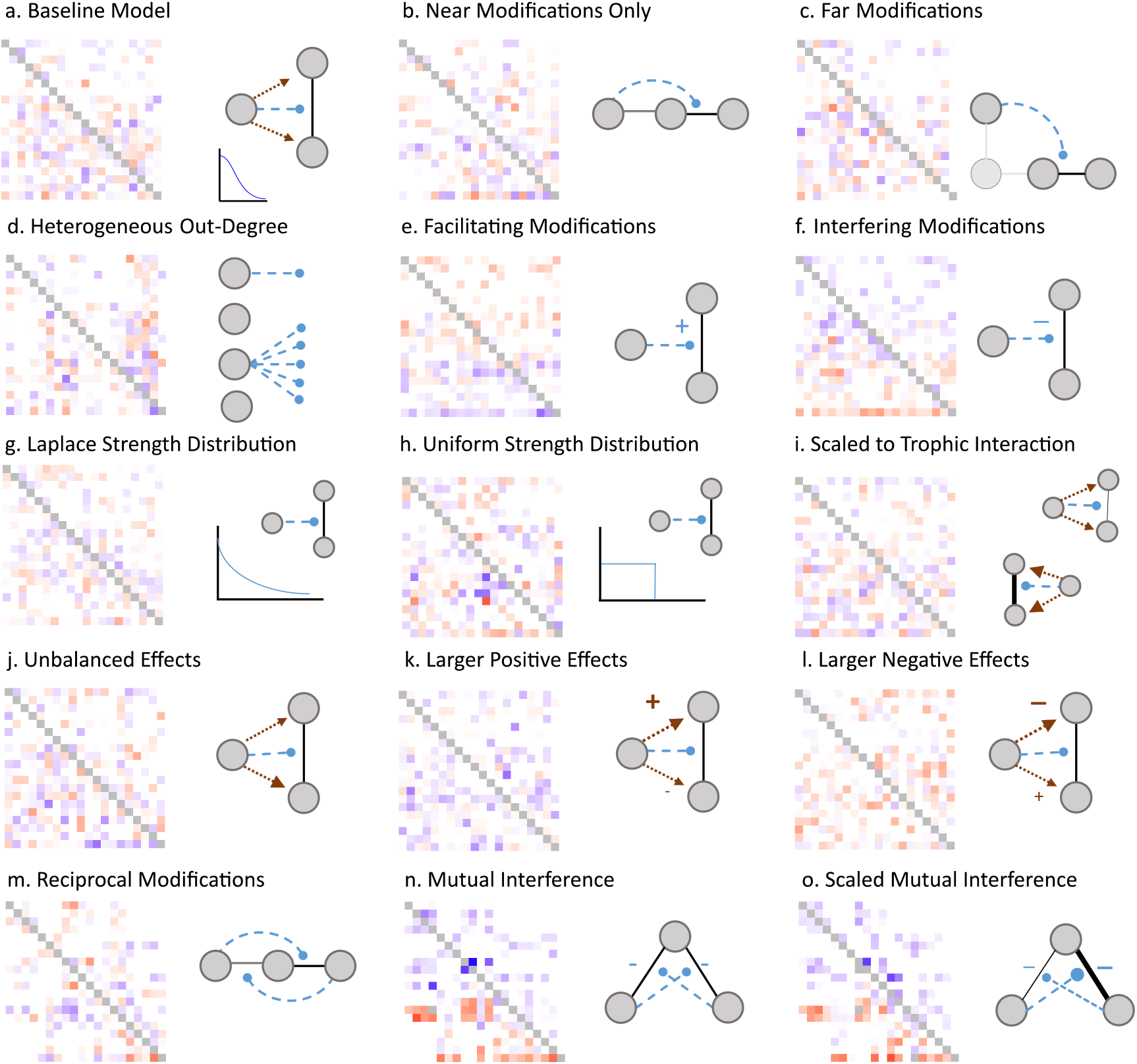
Illustration of the TIMs distribution models used in this study with representative non-trophic effect matrices. Cartoons illustrate the distinctive properties of each model compared to the baseline model (a) that introduced interaction modifications such that each potential modification was equally and independently likely to occur, where the strength of the resultant non-trophic effects were normally distributed and equal on both interactors. See main text and Table S1 for a description of each model.

All TIM models included a dependence on the underlying trophic network, but this interaction modification topology was varied in several of the models. Two models introduced a simple dependence on the trophic distance between the modifier species and the trophic interactors models, testing the effect of interaction localisation within networks. Model b) required that the modifier was trophically connected to at least one of the interactors, while model c) excluded such cases. Model d) tested the effect of certain species exerting a disproportionate number of modifications, representing topologically the impact of certain ‘ecological engineer’ species exerting a dominant effect on the interaction within a network. This was carried out by assigning each species an individual probability drawn from a beta-distribution of being the modifier of each generated interaction modification, thereby generating a more greatly heterogeneous TIM out-degree distribution.

The distribution of TIM strength (the *c_ijk_* parameter in our framework) were varied in two distinct ways. Firstly, the sign of the modification was varied, introducing either exclusively facilitating (model e) or interfering (model f) TIMs. Note that the balance between individual resultant positive and negative NTEs remained constant. Secondly, distributions with different variance of TIM strength were tested. Model g) used a leptokurtic (‘fat-tailed’) Laplace distribution such that there was a greater number of high strength modifications, while maintaining the same mean magnitude. By contrast, model h) used a uniform distribution for *c_ijk_*.

Modifications of stronger trophic interactions are likely to have a greater overall impact. To examine this effect, model i) scaled the strength of the NTEs to the mean strength of the trophic interaction being modified. For this, each *c_ijk_* parameter was drawn from a uniform distribution determined by the relevant elements of the trophic interaction matrix, 𝒰 (−*γ*, *γ*), where 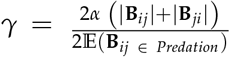. Here the denominator term maintains the overall mean value of the non-trophic effects as half that of the trophic interactions.

The baseline model assumes that each interaction modification induces equally strong positive and negative NTEs, but the balance is unlikely to be equal. In model k) resultant positive NTEs were set at three times the strength of negative NTEs, while in model l) the negative NTEs were three times larger. In model j) whether the positive or negative NTE was larger was randomised to separately test for unbalanced effects on species pairs.

A final set of three models introduced internal structure to interaction modifications. Model m) introduced TIMs only in tightly reciprocal pairs to represent tight clustering of interaction modifications within particular subgroups of species. In this model, sets of three trophically linked species include two reciprocal modifications - a species that modifies an interaction between species x and species y, would in turn have its interaction with x modified by y.

Model n) represents the specific but widespread case of interaction modifications caused by foraging choices between two resources, where each resource reduces the consumption of the other by the shared consumer. This ‘mutual interference’ model therefore includes properties of trophic topology (only trophically nearby modifications are included), sign (only interfering modifications), and internal structure (TIMs are distributed in pairs). Model o) introduces an additional constraint that the size of the NTEs are limited by the size of the trophic interaction being modified, preventing the presence of the additional resource causing a net negative effect on the consumer. This was modelled by drawing *c_ijk_* values from a uniform distribution bounded by the corresponding positive trophic term: *𝒰* (−**B**_*jk*_, 0).

### Properties of interaction matrices

We applied each of the 16 distribution models (including the unstructured NTE model) at each TIM density across each of the 100 underlying trophic networks for a total set of 11 200 communities. For each community we calculated the following structural properties of the resultant interaction matrices: 1) mean interaction strength (*μ*) of **A**, 2) connectance (fraction non-zero entries) of **A** and **C**, 3) variance (V) of the off-diagonal elements of **A** and of **C**, 4) degree heterogeneity of **A** as the variance of the normalised in and out-degree distribution, 5) correlation (*ρ*) of the pairwise elements of 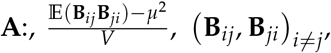 covariance between **B** and **C**, 7) row structure *ζ_row_* as the variance in mean interaction magnitude across rows 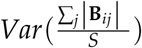 and like-wise: 8) column structure *ζ_col_*, 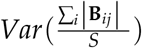. We compared the mean value of each of the above structural properties across the 100 trophic networks for each of the TIM distribution models to the baseline TIM model at a representative TIM density of 10 using a linear model.

We calculated the local asymptotic stability and the reactivity for each community. Local asymptotic stability is determined by the sign of the real part of the leading (dominant) eigen-value of **A**, 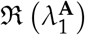, under the assumption that the community is at an equilibrium. If negative, the system will eventually return to the original equilibrium after a small perturbation and is considered locally stable. The diagonal elements of **A** specify the self-regulation of each species. With sufficient self-regulation any community can be stabilised. Although the distribution of self-regulation effects can have important consequences (Barabás *et al.* 2017) we follow previous work (Allesina *et al.* 2015, Jacquet *et al.* 2016) and set all diagonal elements of **A** to zero to focus on the impact of the inter-specific interactions. Without self-regulation, 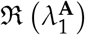 will always be positive but can be interpreted as how far a system is from stability, i.e. how much self-regulation would be necessary to stabilise the system. Hence, although local stability is a binary property, we use ‘less stable’ to refer to a system farther from stability (a larger 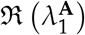).

The immediate response of a system to perturbation is described by its reactivity, the maximum instantaneous amplification of a small perturbation (Neubert and Caswell 1997, Tang and Allesina 2014). This is computed as the leading eigenvalue of the Hermitian part of the community matrix: 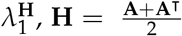. Since **H** is symmetric, its eigenvalues are real.

To assess the extent to which the communities were non-randomly structured we tested the performance of two analytic criteria for local stability derived from generalisations of the circular law of random matrix theory (May 1972, Tang *et al.* 2014, Allesina and Tang 2015). We refer to these as the May criterion: 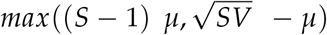 and the Tang *et al.* criterion: 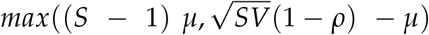. Here, *S* is the number of species (the size of the matrix), *V* is the variance in interaction strength, *μ* is the mean interaction strength and *ρ* is the pair-wise correlation between interaction terms, 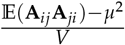. The first term of each criteria specifies the eigenvalue derived from the expected row-sum (Allesina and Tang 2012). In most trophic networks (where *μ* ≪ 0), this term is usually the smaller and can be safely ignored, but this is not necessarily true with the addition of non-trophic effects. The second term estimates the radius of the ellipse containing the eigenvalues along the real axis, and hence the likely position of the right-most (dominant) eigenvalue, under the assumptions of random matrix theory: that the matrix is large and the entries are independent and identically distributed (Tao 2012, Allesina and Tang 2015).

### Empirical trophic networks

The stability analysis was repeated for the five largest empirical trophic networks in the set compiled by Jacquet *et al.* (2016). These are models of the trophic interactions in marine fisheries that had been parameterised by the EcoPath modelling approach (Christensen and Pauly 1992) and converted to Jacobian matrices following the method of de Ruiter *et al.* (1995). These networks were: Chesapeake Bay (n = 41, links = 167, Christensen *et al.* 2009), Mid Atlantic Bight (n = 51, links = 515, Okey and Pugliese 2001), Moorea Barrier Reef (n = 39, links = 267, Arias-González *et al.* 1997), mid-1990s Newfoundland Grand Banks (n = 48, links = 525, Heymans and Pitcher 2002) and Tampa Bay (n = 48, links = 340, Walters *et al.* 2005).

Interaction modifications were introduced at densities of 1, 5, 10, 20, 30 and 40 TIMs per species with 100 replicates of each TIM distribution model at each density for each of the trophic networks for a total set of 48 000 communities. The mean strength of the interaction modifications (*α*) introduced to each trophic network was set at half the mean strength of the positive trophic interactions (range 0.017-0.18) to be comparable with the artificial networks.

## Results and Discussion

In the artificial networks, as the density of TIMs included in the model was increased, local stability always decreased while reactivity always increased (Figure 4a, b). Within this overall pattern, a number of TIM distributions led to markedly different responses, discussed below. Although the remainder of the TIM models had impacts on stability that were not meaningfully different to random NTEs, there was a small but discernible split in their reactivity response (Figure S1b). Models that led to NTEs being focussed, clumped or unbalanced (Larger Negative NTEs, Laplace strength distribution, Nearby TIMs, Unbalanced NTEs, Heterogeneous out-degree and Reciprocal modifications models) led to a higher reactivity than those that had a more even effect distribution (Baseline, Scaled, Random NTE, Far TIMs, Uniform strength distribution).

**Figure 4.**
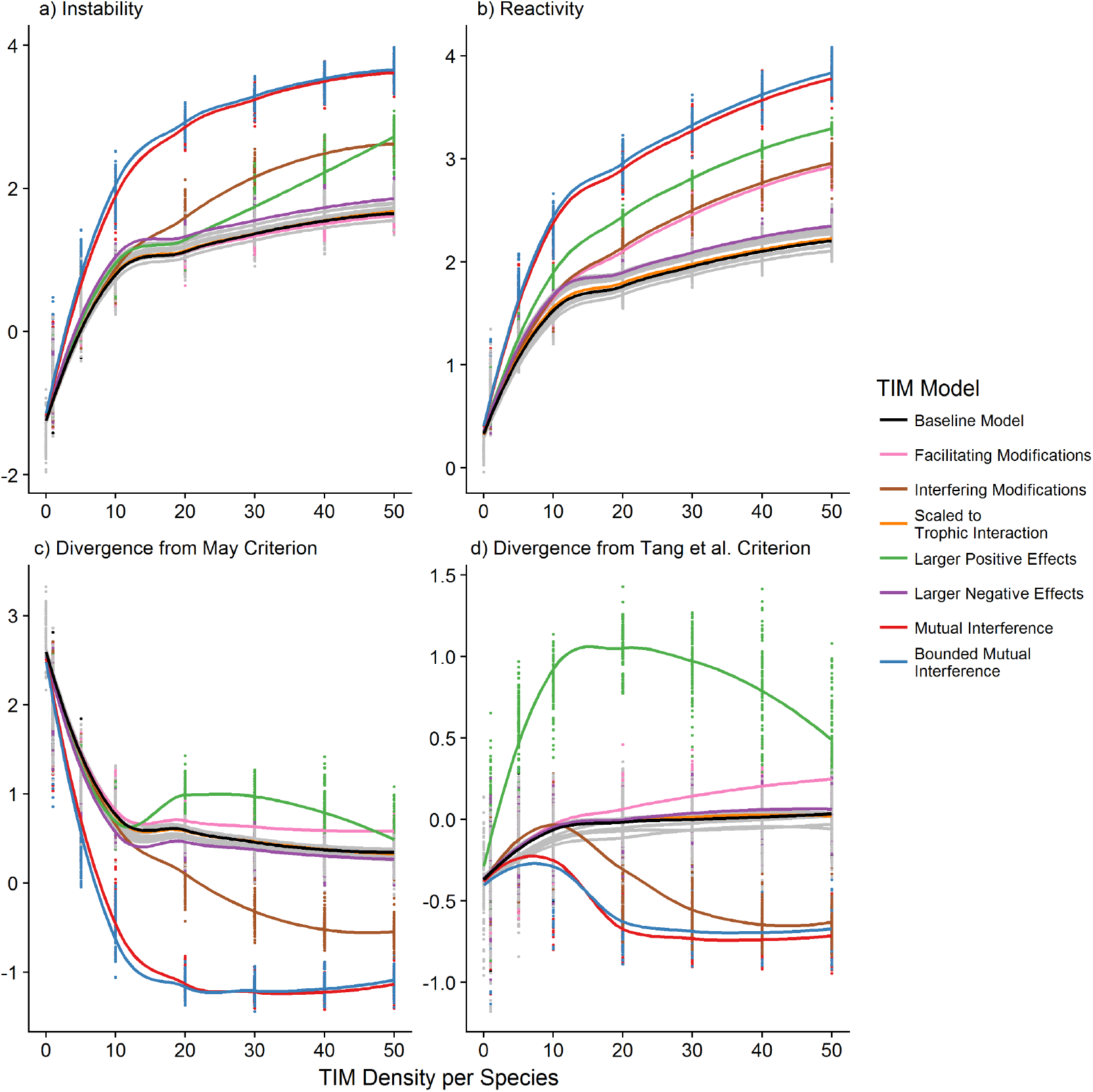
Effect of increasing density of TIMs on (a) instability 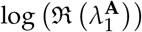, the degree of self-regulation necessary for local asymptotic stability, (b) system reactivity 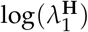, (c) the log-ratio of the May stability criterion and the observed stability and (d) the log-ratio of the Tang *et al.* stability criterion and the observed stability. TIM distribution models with distinctive responses are highlighted, the remainder are coloured grey and are plotted in Figure S1. Loess fitted lines have been added to highlight differences.

The two mutual interference models, representing cases where two resources that share a consumer both negatively affect the consumption of the other (Figure 3.n-o), had the greatest impact on stability and reactivity. At higher levels of TIM density, interfering modifications (which weaken a trophic interaction and are beneficial for the resource and detrimental for the consumer, at least in the short term, Figure 3.f) caused greater instability and reactivity than other models. Facilitating modifications (Figure 3.e) did not cause a distinct impact on stability, but the impact on reactivity closely matched the interfering modification distribution. The distribution with larger positive NTEs (Figure 3.k) also caused distinct effects, reducing stability at high TIM densities but increasing reactivity faster than baseline models at all TIM densities. Results for further analyses where trophic interactions negatively affected resources more than consumers benefitted followed broadly the same pattern, with the exception that the model with larger positive NTEs had reduced impact as the overall mean interaction strength remained negative (Figure S2 & S3).

Many of the TIM distributions we tested showed a reduced divergence of the true stability from that estimated by the stability criteria of both May and Tang *et al.* (Figure 4c and 4d). This suggests that the TIMs are moving the systems closer to fulfilling the assumptions of random matrix theory under which the criteria are derived, namely that individual interactions are independently and identically distributed (Tao 2012, Allesina and Tang 2015), despite the observable structure being introduced (S.I. Table S2). However, the same TIM models that significantly impacted stability, increase the divergence between the true stability of a system and that estimated by either stability criteria, showing that the TIMs are having consequences which are not captured by the mean, variance and correlation of the matrix elements. All three interference TIM models led to communities that were more stable than expected by the criteria while larger positive NTEs and facilitating TIMs were less stable. Simple, but plausible, distributions of TIMs can therefore be seen to push communities either closer or further (in either direction) from expectations based on the study of random matrices (James *et al.* 2015) than may be expected from analyses of trophic interactions alone.

The structural feature of the resultant matrices that can best explain stability across the set of communities was *ρ*, the correlation between pairwise elements of the overall community matrix **A**, with an *r*^2^ of 0.884 over all the generated communities (Figure S4). This can also be observed in the marked superiority of the Tang *et al.* criterion (which includes *ρ*) over the May criterion (Figure 4c-d), although pairwise correlation was a less good predictor of stability at weak (near zero) levels of correlation (Figure S4a).

### Mutual Interference TIMs

The mutual interference model of TIMs is strongly destabilising, despite the lower variance in overall interaction strength (since negative non-trophic interactions overlap with positive trophic interaction terms), which would normally be expected to increase stability (May 1972). This effect is driven largely by emergent pairwise mutualism, long-recognised as destabilising for interaction matrices (May 1973). The NTEs induced by these TIMs are very efficient at breaking down the negative correlation between pairwise elements (Figure S4b), inducing mutualistic effects between resources that share a consumer. However, the maintained divergence from the Tang *et al.* criteria, which includes pairwise correlation, suggests additional higher-level structural contributions.

The mutual interference models result in a high variance in the elements of the NTE matrix **C** since they tend to focus negative and positive NTEs on to distinct groups of species (the high-level consumers and low-level resources). However, since in this model resources exert negative NTEs upon their consumers, matrices **B** and **C** have a low covariance, and the resultant variance of **A** is lower than that derived from other NTE distributions. The importance of the sign structure of interference can also be seen in the comparative lack of distinction of the tightly reciprocal interaction modification distribution (Fig 3m) compared to random NTE distributions - it is the specific sign patterning of the links that drives the change to the dynamics, not the topological clustering of the modifications.

Reciprocal negative effects between consumers can be generated by a range of mechanisms, including predator satiation (Jeschke *et al.* 2004), adaptive foraging (Abrams 2010) and associational defence (Barbosa *et al.* 2009). These effects are widespread (Bruno *et al.* 2003) and are regularly included in general models of population dynamics (Delmas *et al.* 2017) through multi-species functional responses (Koen-Alonso 2007), yet the resultant dynamic links are rarely considered in network based analyses. The divergence with conclusions drawn from small community modules, where switching is generally considered stabilising (Murdoch and Oaten 1975, May 1977, Valdovinos *et al.* 2010), reinforces the need to consider dynamics across scales and mechanisms.

### Sign Effects of TIMs

Interfering TIMs, which are beneficial for resources and detrimental for consumers, have a strongly destabilising effect at high TIM densities. The interfering TIM model differs from the facilitating TIM model only in the sign patterning of the NTEs, yet it has greatly different effects on stability because of the relationship with the underlying trophic interactions. Under these distributions, species at either end of food chains are only susceptible to one sign of NTE, resulting in the banded sign pattern observable in Figure 3e and 3f. Since there are distinct patterns in the underlying trophic interaction distributions (Figure 2) the covariance between matrices **B** and **C** is higher for facilitating modifications than interfering (Table S2). Hence, a large number of interfering interaction modifications tends to lead to weaker overall pairwise exploitative relationships, despite our model keeping underlying trophic interaction strength fixed. This reduces the variance and pairwise correlation, *ρ*, of **A** faster than randomly signed TIMs (e.g. with a TIM density of 10, *ρ* = −0.55 compared to −0.61, Table S2). Conversely, facilitating modifications break down the pairwise correlation slightly slower (*ρ* = −0.62, at TIM density of 10) than random interactions.

Both facilitating and interfering TIM distributions had similar effects on reactivity (Figure 4b). Reactivity (Neubert and Caswell 1997, Tang and Allesina 2014) is dependent on the eigenvalues of 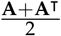, effectively a matrix composed of mean pairwise interaction strengths. Both facilitating and interfering TIM distributions result in strong row patterning of this matrix. In biological terms, certain species at either end of food chains accrue consistently stronger interactions, generating reactivity. While NTEs from TIMs are not directly reciprocal, if either interfering or facilitating modifications are more common, consistent patterns can develop across trophic levels.

In the case where positive NTEs were larger than negative NTEs, changes in stability were driven by the increasingly positive mean interaction strength. This is highlighted by the stability criterion estimate being determined by the expected row sum component, (*S −* 1) *μ*, of the criterion for all but cases with the fewest TIMs. There was no particular impact of the imbalance itself – both the unbalanced NTE model and the model where negative NTEs were consistently larger were not notably different from the random NTE case.

At present, the empirical balance between facilitating and interfering modifications and their distribution throughout ecological communities is effectively unknown, as is the balance between resultant positive and negative consequences for interactors. While the all or nothing cases discussed here represent extreme cases, they show that this data will be essential in determining the impact of non-trophic effects on dynamics.

### TIMs and Row Structuring

It has recently been suggested that row-structuring (where the rows of interaction matrices have markedly different means) is a significant contributor to the stability of empirical networks (Jacquet *et al.* 2016), although the effects may be greater on the rate of return to equilibrium than the sign of the eigenvalue (Gibbs *et al.* 2017). Row structuring in trophic networks is a consequence of the consistent dependence of population level interaction strength on interactor density. TIMs may cause row and column structuring by two distinct mechanisms. Firstly, dominant species that take part in strong interactions may be expected to be the receiver of strong NTEs caused by interaction modifications – even a small change in a large interaction could be expected to have a large overall effect on the flow of energy through a community as a whole. Secondly, species that induce a disproportionate number of interaction modifications (ecological engineers (Jones *et al.* 1994, Sanders *et al.* 2014)) have the potential to introduce considerable column-structuring effects. Such ecological engineers causing a large number of interaction modifications may not necessarily be those involved in the strongest trophic interactions. NTEs exerted by such species may serve to break down or replace existing column structure. However, our results suggest that high variation in the number of TIMs each species exerts (out-degree heterogeneity, Figure 3d), representing such a scenario, does not affect stability differently to random NTEs in the artificial trophic networks.

### Empirical Trophic Networks

When we repeated our analysis for five empirical marine fishery trophic networks in the set compiled by Jacquet *et al.* (2016) we found distinct results (Figure 5). In contrast to the artificial networks, models where NTEs scaled with the underlying trophic interactions had by far the most significant effects on dynamics of the empirical networks. This occurred despite the mean magnitude of individual NTEs being kept constant between methods of introducing TIMs. In 4 out of 5 cases mutual interference led to more instability, although in the one case (the ‘Newfoundland’ network) the effect was opposite – the unscaled mutual interference model was stabilising. In another case (the ‘Mid Atlantic Bight’ network), many of the TIM models led to a small increase in stability, with the largest effect from facilitating modifications.

**Figure 5.**
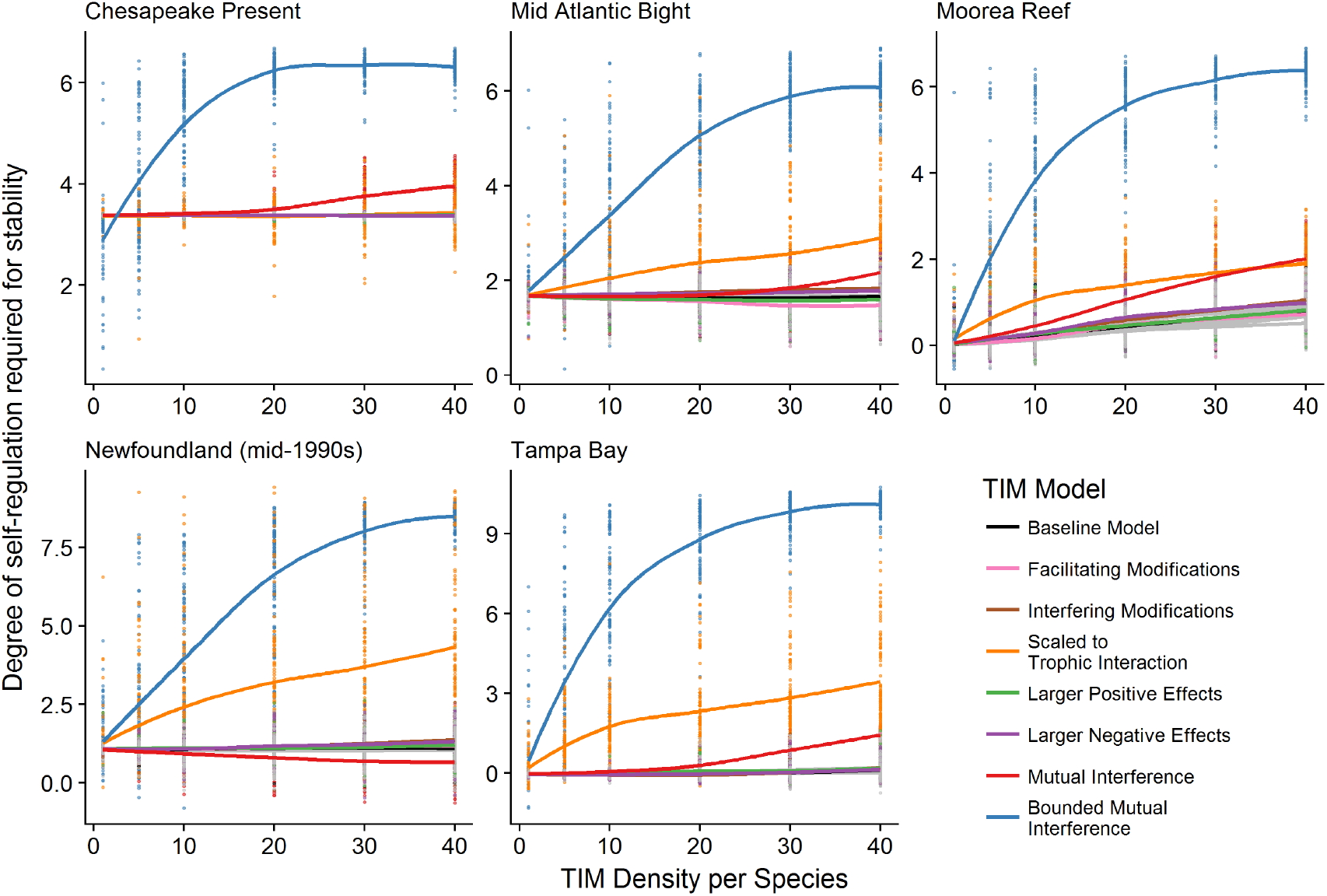
Stability responses of five empirically parameterised food webs to increasing density of TIMs. TIM distribution models with distinctive responses are highlighted, the remainder are coloured grey and are plotted in Figure S5. Loess fitted lines have been added.

The empirical networks studied here differ significantly from randomly generated networks (see Jacquet *et al.* (2016) for a full discussion). They include both significant row-structuring and an approximately log-normal interaction trophic strength distribution. A minority of very strong, mostly negative, interactions represent large transfers of biomass between dominant species in the community (SI 5), resulting in a highly leptokurtic distribution of trophic interaction strengths and a very low correlation coefficient (range *ρ* = −0.014:−0.001). In such a case, unless the TIMs are scaled with the underlying trophic interactions, most interaction modification distributions in our test set effectively only add noise. Consequently, the introduction of TIMs hardly changed the overall pairwise correlation - the largest value of *ρ* across all TIM iterations was just −0.016. The NTEs introduced by the scaled models matched the highly leptokurtic distribution of the trophic interactions leading to an increased variance in NTEs. However, even the increase in variance of the ‘scaled’ distributions was several orders of magnitude smaller than the variance in the underlying trophic interactions (SI 6). Hence, the stability criteria were almost completely unresponsive to the introduction of TIMs (Figure S6). Nevertheless, a handful of strong NTE links can be observed to cause great destabilisation, while in most cases the comparatively weak links introduced by TIMs from constrained, unscaled, distributions had very little effect on stability.

## Conclusion

Our results show that trophic interaction modifications have the potential to cause significant disruptive effects to the pattern of interactions within ecological communities. The inter-relationship between types of interaction is critical to understanding their consequences, reiterating the importance of the dependencies between superimposed interaction networks (Kéfi *et al.* 2016, Pilosof *et al.* 2017). TIMs can influence the stability of systems through a number of distinct mechanisms beyond introducing additional connectance. They can shift the average interaction sign, change pairwise correlation coefficients, introduce additional row structure and change the interaction strength distribution. Furthermore, given the potential for interaction modifications to short-circuit established trophic interaction motifs such as tri-trophic cascades, the distribution of interaction motifs in ecological communities (Milo 2002, Stouffer *et al.* 2007) may need to be re-examined to incorporate non-trophic effects.

Empirical data on the distribution of interaction modifications in real communities will be essential to discern their true effects. The distribution of non-trophic effects, including those caused by interaction modifications, in real communities is at present essentially unknown beyond a limited number of inter-tidal communities (Kéfi *et al.* 2015, Sander *et al.* 2015). The fraction of interspecific interactions driven by interaction modifications is unknown, but likely to be large (Abrams 1983, Werner and Peacor 2003). The set of models used here attempts to map some of the properties real distributions of interaction modifications could have and identify features pertinent to dynamics. Thus, they provide a stepping stone between analytical random matrix approaches (Bairey *et al.* 2016) and empirically parameterised systems.

Ideally, empirical data will need to include information about both the topology and strength of the non-trophic, which is possible, but challenging, to acquire (Mayfield and Stouffer 2017, Terry *et al.* 2017). This could lead to the identification of regular patterns across ecosystems in the features identified here as pertinent to community dynamics. Whilst it is unlikely that there will be strong mechanistic drivers of non-trophic network structure equivalent to the role of body-size within trophic interaction networks (Petchey *et al.* 2008, Brose 2010, Pawar *et al.* 2012), there is nevertheless room for a great improvement in our phenomenological understanding of the distribution of interactions modifications.

Studies focussed exclusively on trophic networks are missing a large portion of the dynamic interactions occurring in ecological communities. Ultimately, the dynamics of ecological communities are dependent on the strength of interactions not whether they are caused by trophic or non-trophic effects. While this study is limited by both its equilibrium setting and by excluding direct non-trophic interactions (such as competition) and the modifications of direct non-trophic interactions (Grilli *et al.* 2017, Mayfield and Stouffer 2017, Levine *et al.* 2017). Empirical work will be necessary to compare the relative contribution of trophic and no-trophic networks and modifications to them. It is nonetheless clear that trophic interaction modifications can provide a framework to leverage our understanding of trophic networks to make significant inroads into the study of non-trophic effects. The key questions in ecological dynamics cannot be satisfactorily resolved until non-trophic interactions are fully integrated into community ecology. Until the empirical distribution of interaction modifications is understood, suggested resolutions of the stability-complexity paradox may be premature.

## Data and Code Availability

All generated data and R scripts used in the analysis is available on OSF DOI: 10.17605/OSF.IO/6FNAV

## 1: Model Specifications

**Table S1:**
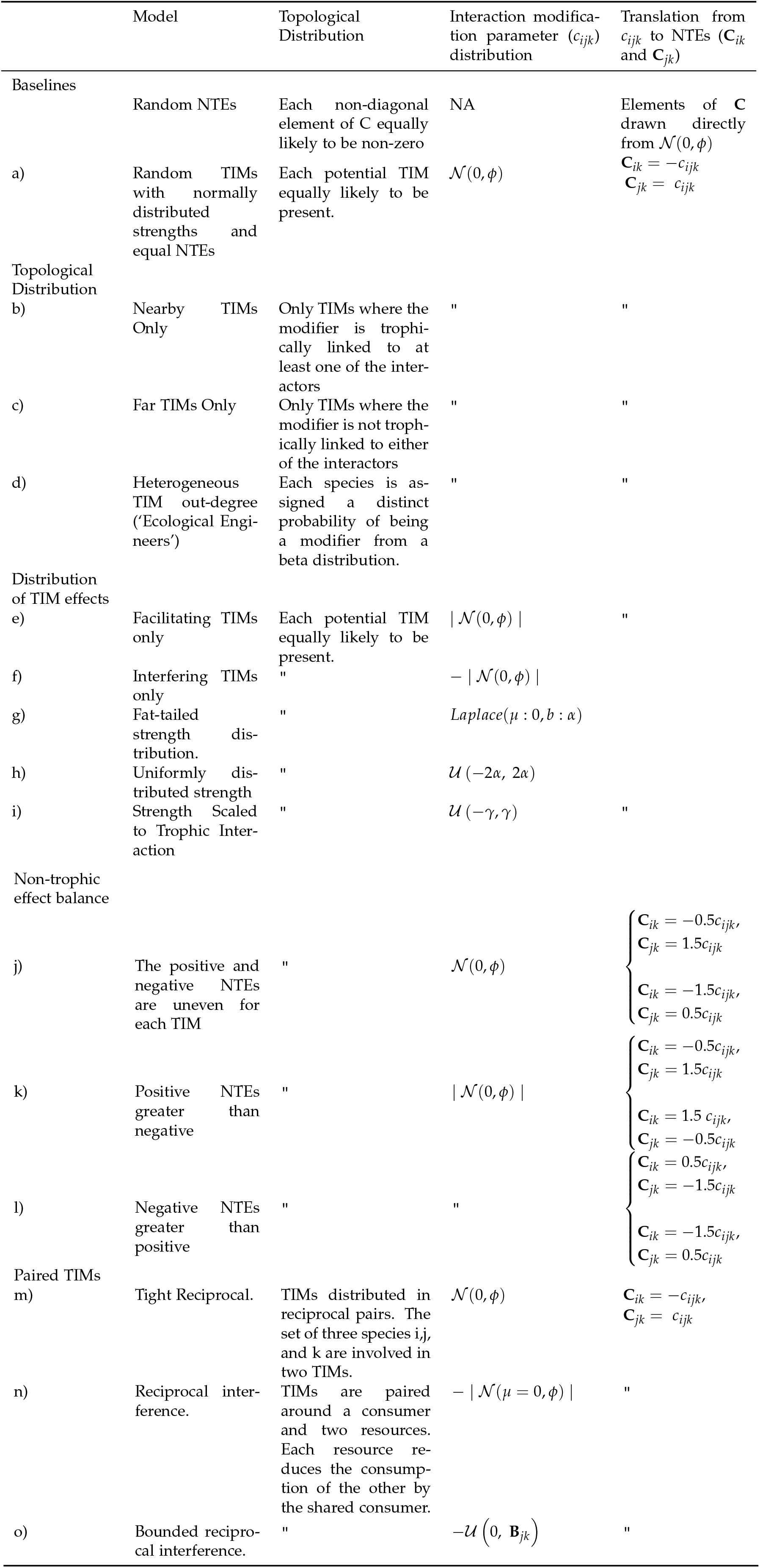
Description of models used to distribute interaction modifications and determine non-trophic effects. Letters match the diagrams in the main text. *α* is the mean magnitude of interaction modification strength, *b_ij_* denotes trophic interaction coefficients, *i* denotes the resource, *j* the consumer and *k* the modifier species of a given interaction modification. Parameter *ϕ* is a standardisation terms used in the Normal distribution to maintain 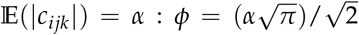 fulfills the same role in the scaled model: 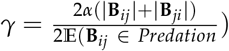

## 2: Structural Properties of Artificial Networks

**Table S2:**
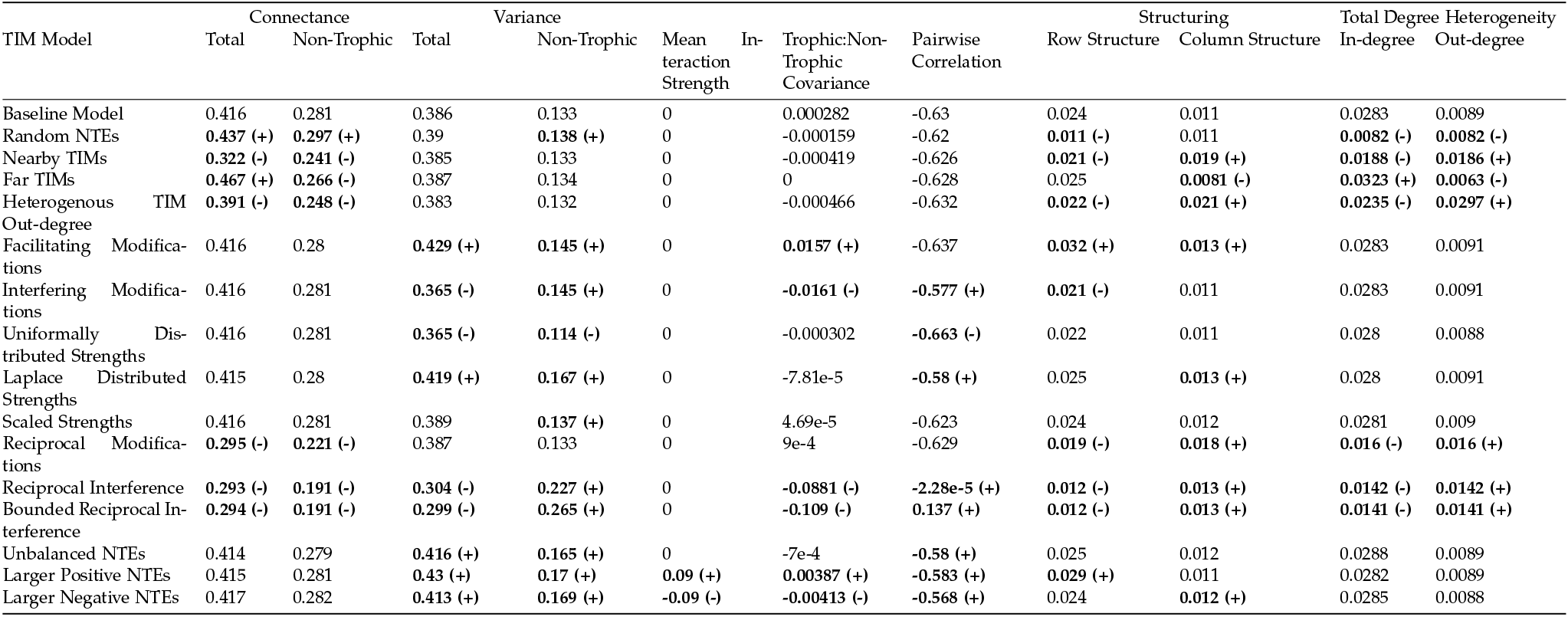
Mean value of structural properties of the communities with TIMs introduced under different models at a TIM density of 10 per species. To aid interpretation those models that differed significantly (linear model, p¡0.05, n=100 per model) from the baseline of equally distributed TIMs with a Normal strength distribution (first row) for a particular structural feature are shown in bold with a (+) or (-) indicating the direction of difference. Exact values of estimated differences, t-values, and p-values are listed in ‘Full Statistical Results.csv’, available at https://osf.io/8zrsm/

## 3: Additional Stability and Reactivity Results

### Non-highlighted TIM models

**Figure S1:**
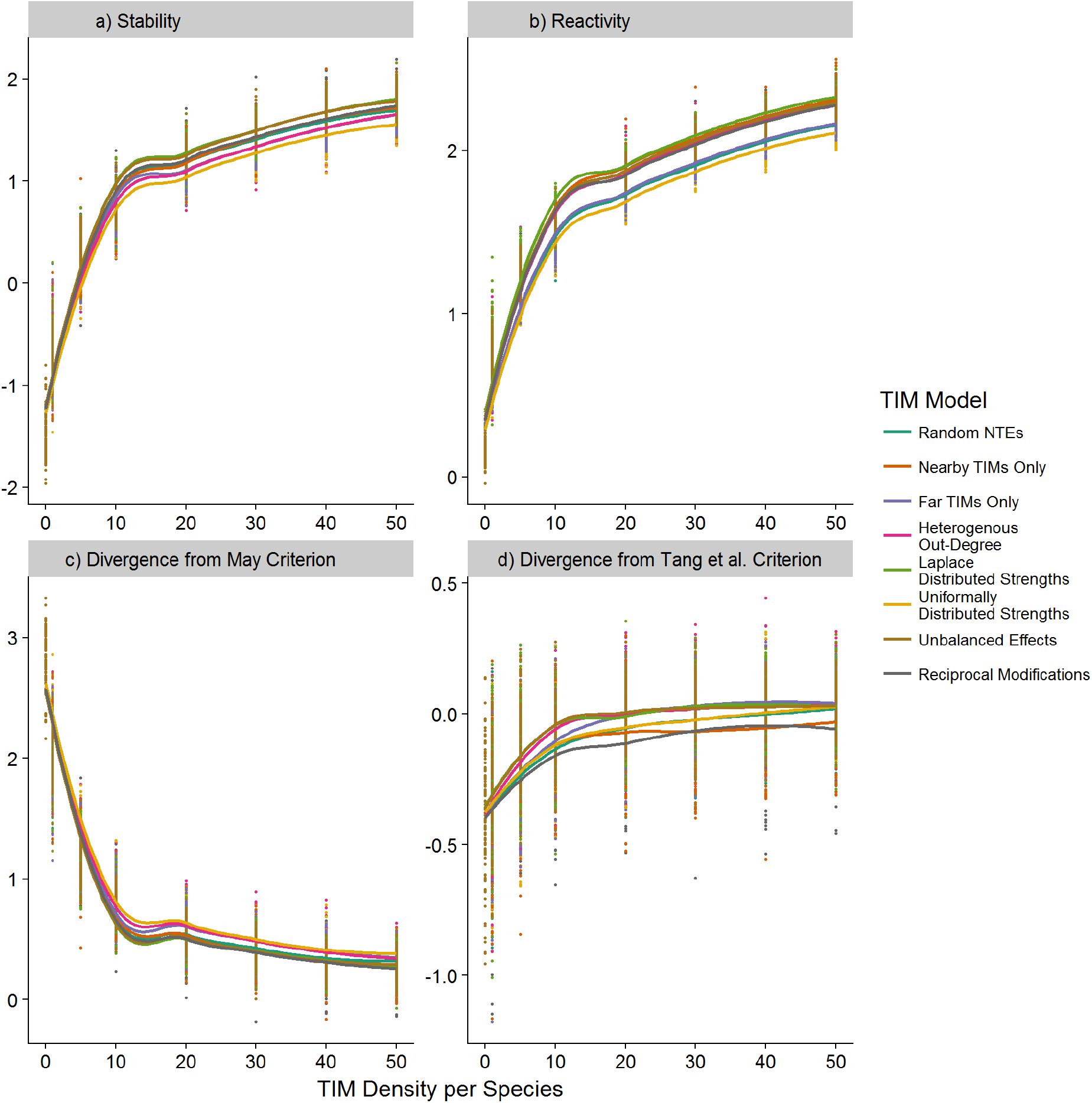
Responses of models that were coloured grey in the main text Figure 4. Effect of increasing density of TIMs on (a) stability 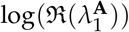, the degree of self-regulation necessary for local asymptotic stability, (b) system reactivity 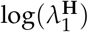, (c) the log-ratio of the May stability criterion and the observed stability and (d) the log-ratio of the Tang *et al.* stability criterion and the observed stability.

**Figure S2:**
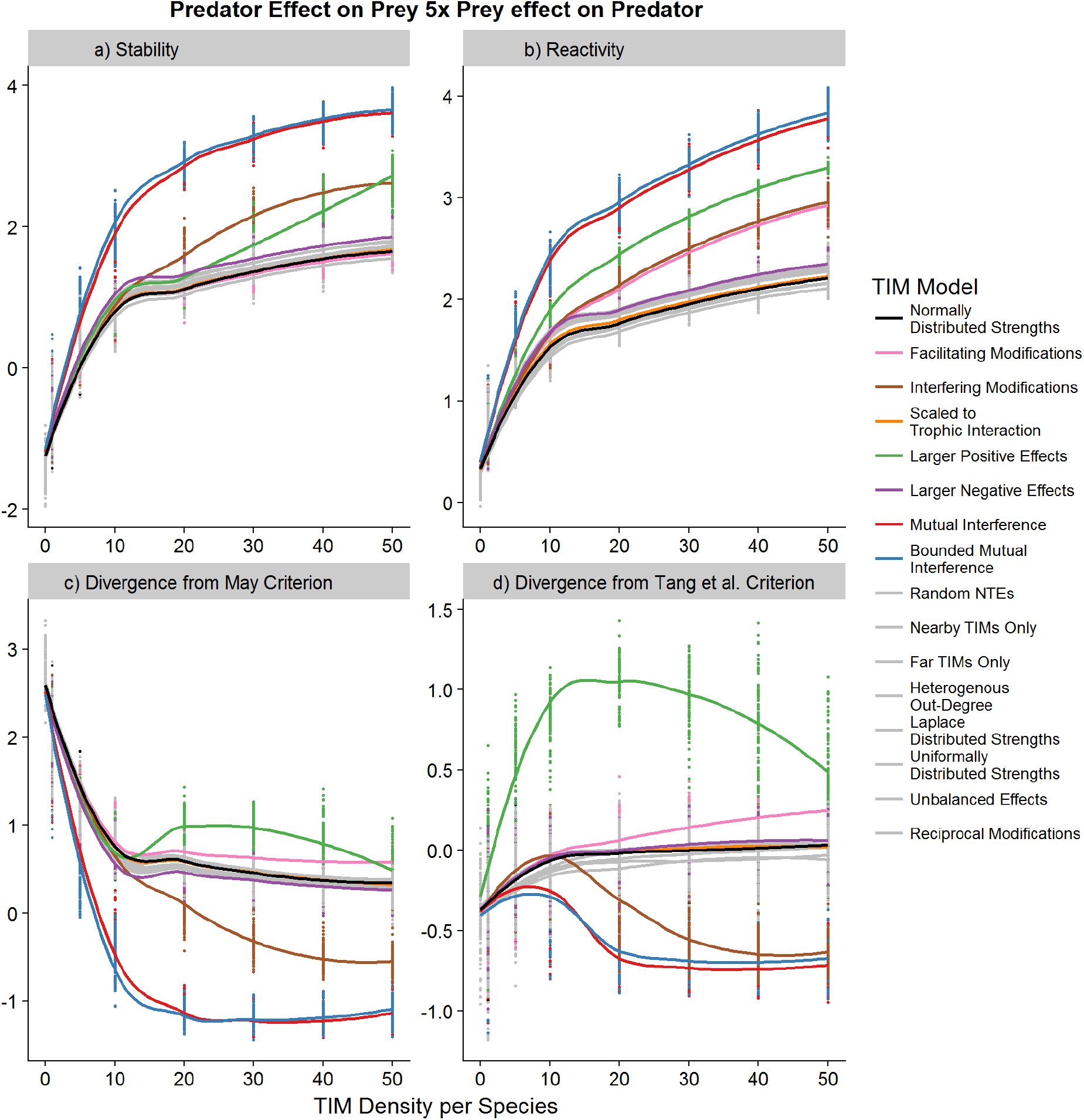
Repeat of results shown in main text Figure 4 with an underlying trophic interaction distribution where resources were negatively affected by consumers on average 5x as much as consumers were affected by resources. Panels show effect of increasing density of TIMs on (a) stability 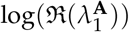, the degree of self-regulation necessary for local asymptotic stability, (b) system reactivity 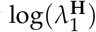, (c) the log-ratio of the May stability criterion and the observed stability and (d) the log-ratio of the Tang *et al.* stability criterion and the observed stability. The impact of TIMs are comparable with the main text results, with the exception that larger positive NTEs (green line) do not lead to destabilisation and facilitation TIMs (pink line) no longer have a distinctive effect.

**Figure S3:**
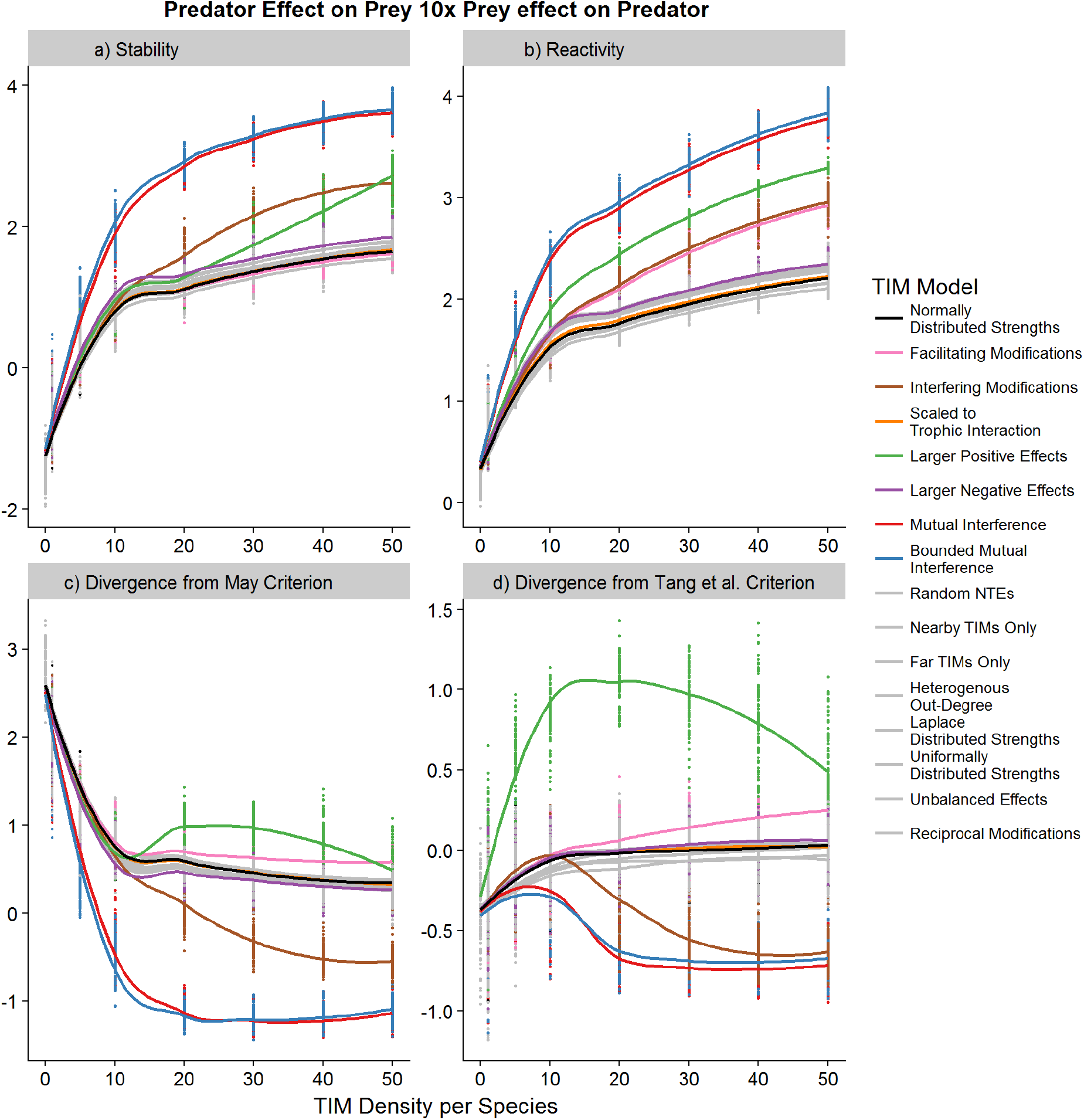
Repeat of results shown in main text Figure 4 with an underlying trophic interaction distribution where resources were negatively affected by consumers on average 10x as much as consumers were affected by resources. Panels show effect of increasing density of TIMs on (a) stability 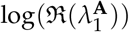, the degree of self-regulation necessary for local asymptotic stability, (b) system reactivity 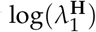, (c) the log-ratio of the May stability criterion and the observed stability and (d) the log-ratio of the Tang *et al.* stability criterion and the observed stability. As with the results shown in Figure S2, the impact of TIMs are comparable with the main text results, with the exception that larger positive NTEs (green line) do not lead to destabilisation and facilitation TIMs (pink line) no longer have a distinctive effect. The grey line that has a less destabilising effect and which maintains the greatest deviation from the Tang et al. Criterion is the Reciprical Modifications model.

**Figure S4:**
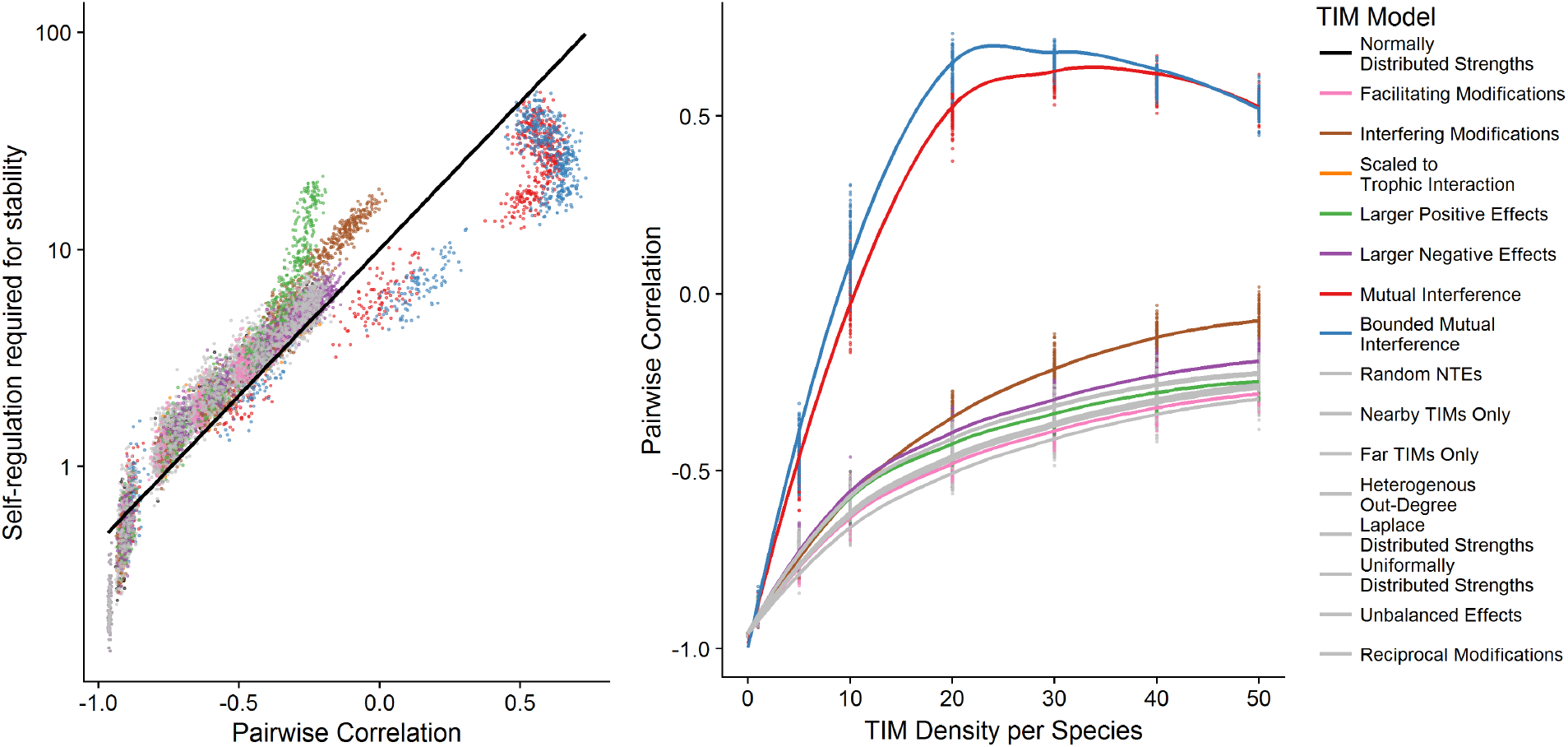
a) Scatter plot across all replicates and models of the relationship between the pairwise correlation between overall interactions and the stability of the system 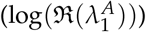 with the artificial trophic networks. The line shows a linear model. It can be seen that the pairwise correlaiton is a very strong predictor of stability. b) shows the impact of TIM density on the pairwise correlation across the different models, with the most diveregence models highlighted. The two mutual interference models most rapidly break down the original negative correlation caused by the specification of the trophic interactions

## 4: Empirical Network Interaction Strength Distribution

Plots of the approximately log-normal distribution of trophic interaction strengths in the five empirical webs, split by whether the interactions were positive or negative.

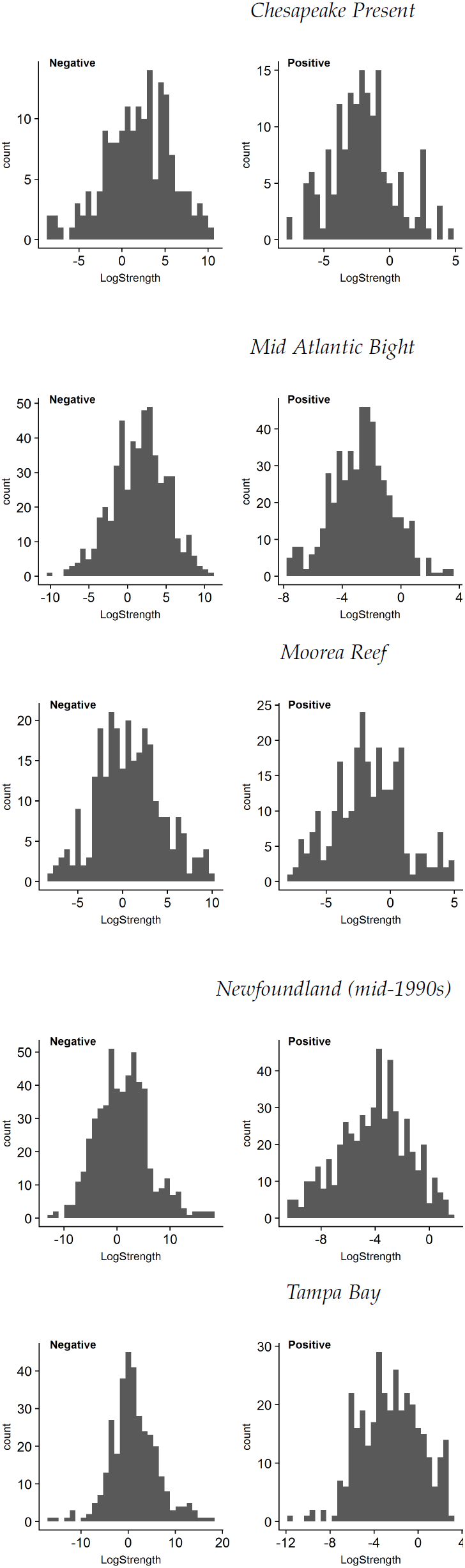

## 5: Structural Properties of Empirical Networks with TIMs

### Chesapeake Present

**Table S3:**
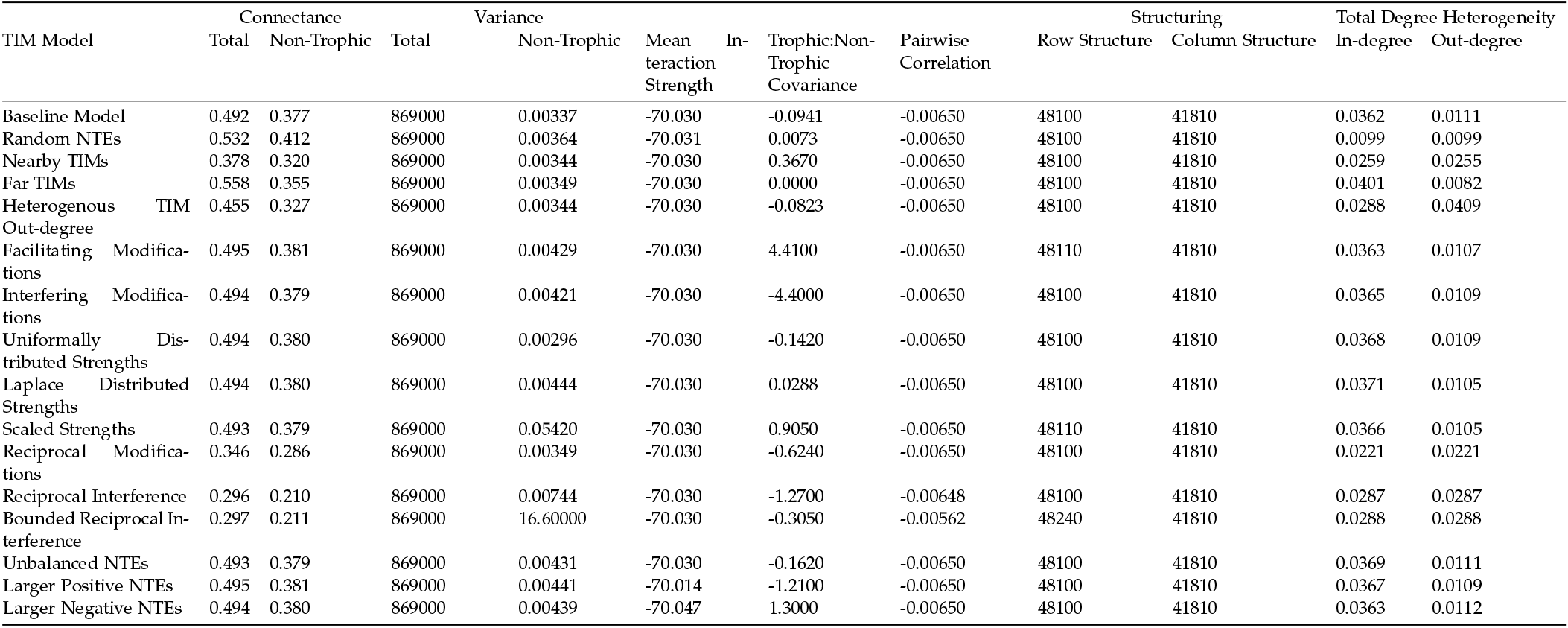
Table showing structural properties of Chesapeake Present network with added TIMs.

### Mid Atlantic Bight

**Table S4:**
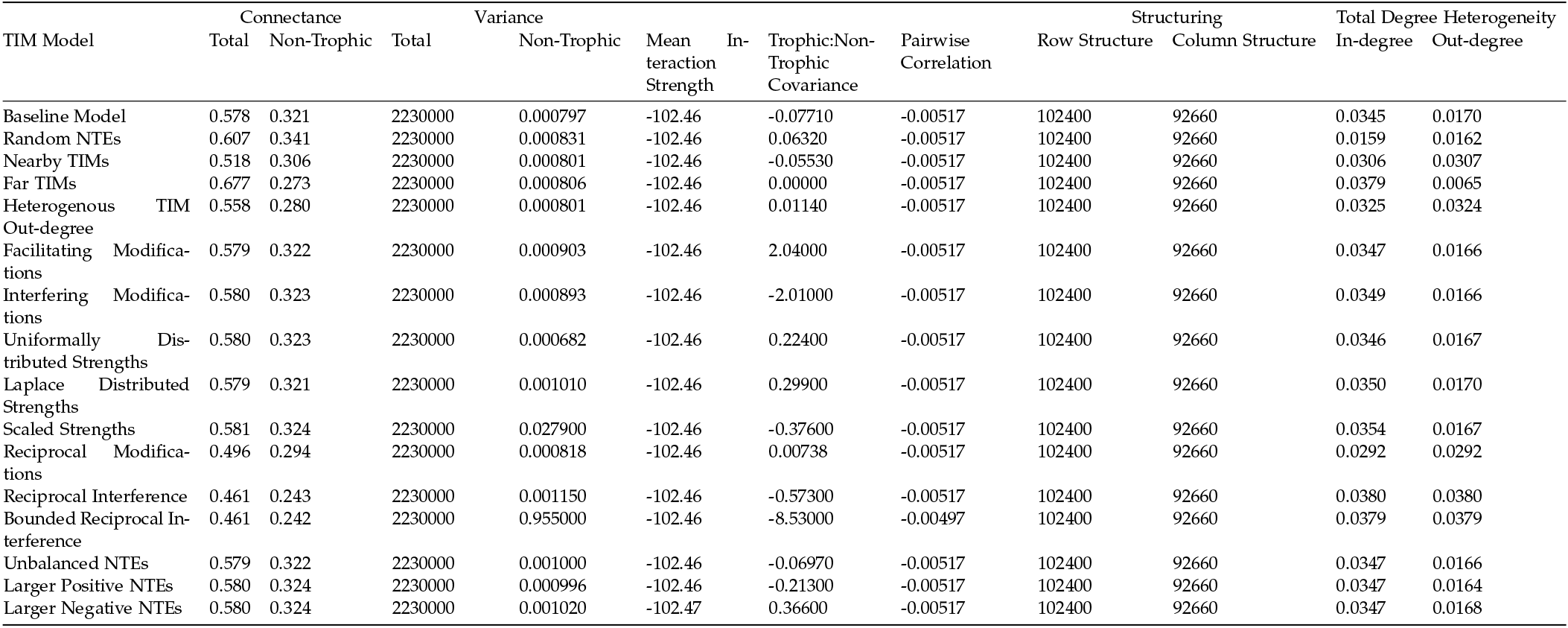
Table showing structural properties of Mid Atlantic Bight network with added TIMs.

### Moorea Reef

**Table S5:**
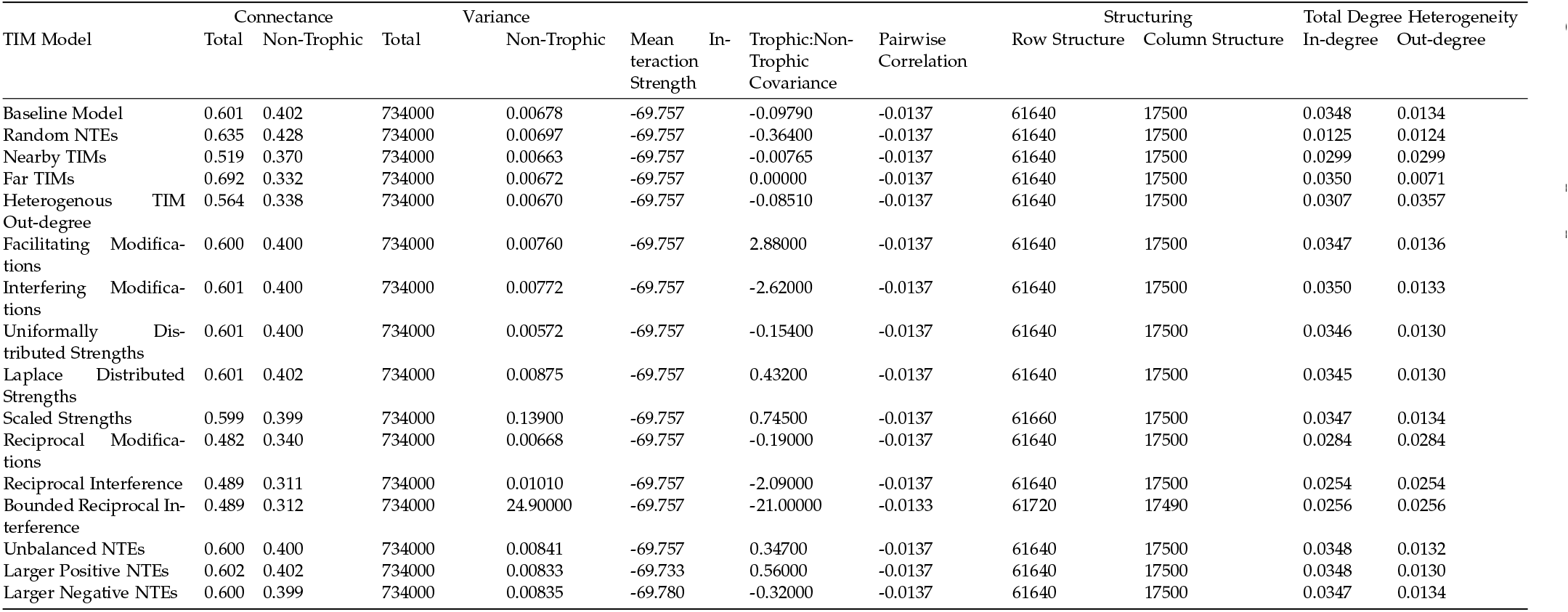
Table showing structural properties of Moorea Reef network with added TIMs.

### Newfoundland (mid-1990s)

**Table S6:**
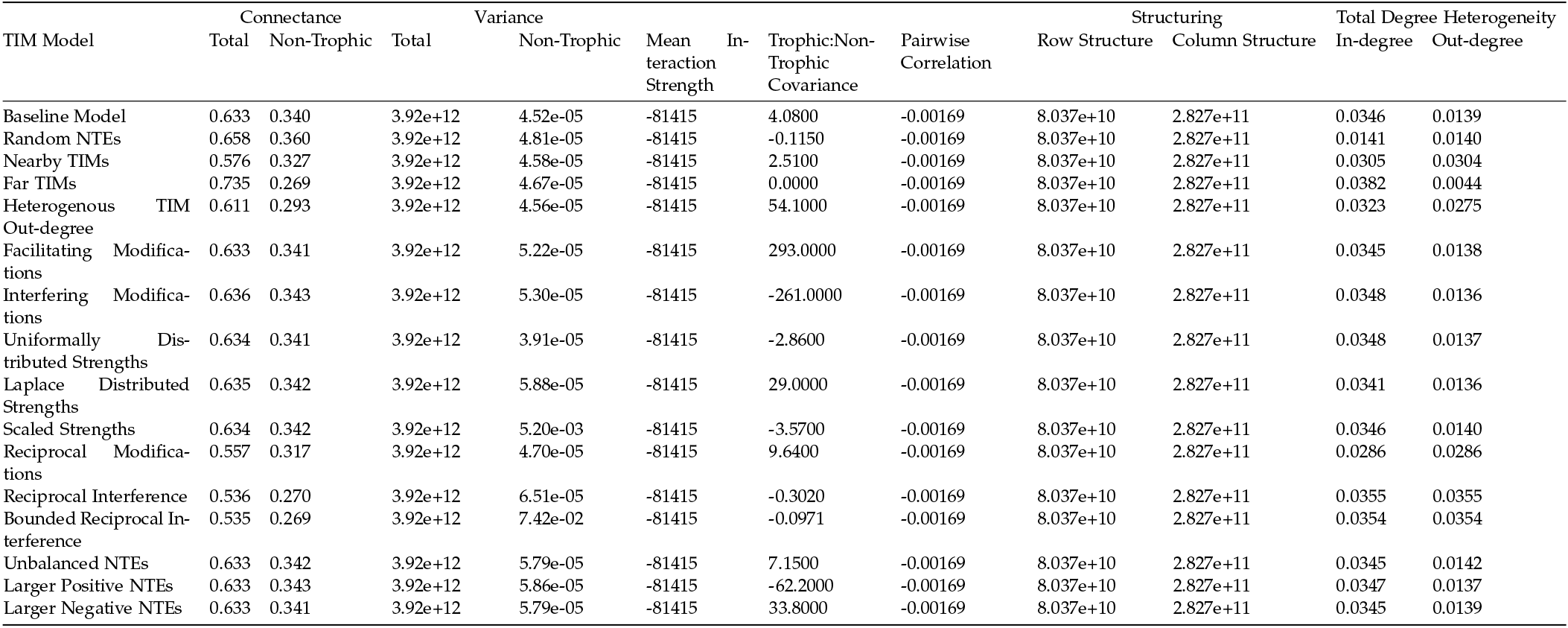
Table showing structural properties of Newfoundland (mid-1990s) network with added TIMs.

### Tampa Bay

**Table S7:**
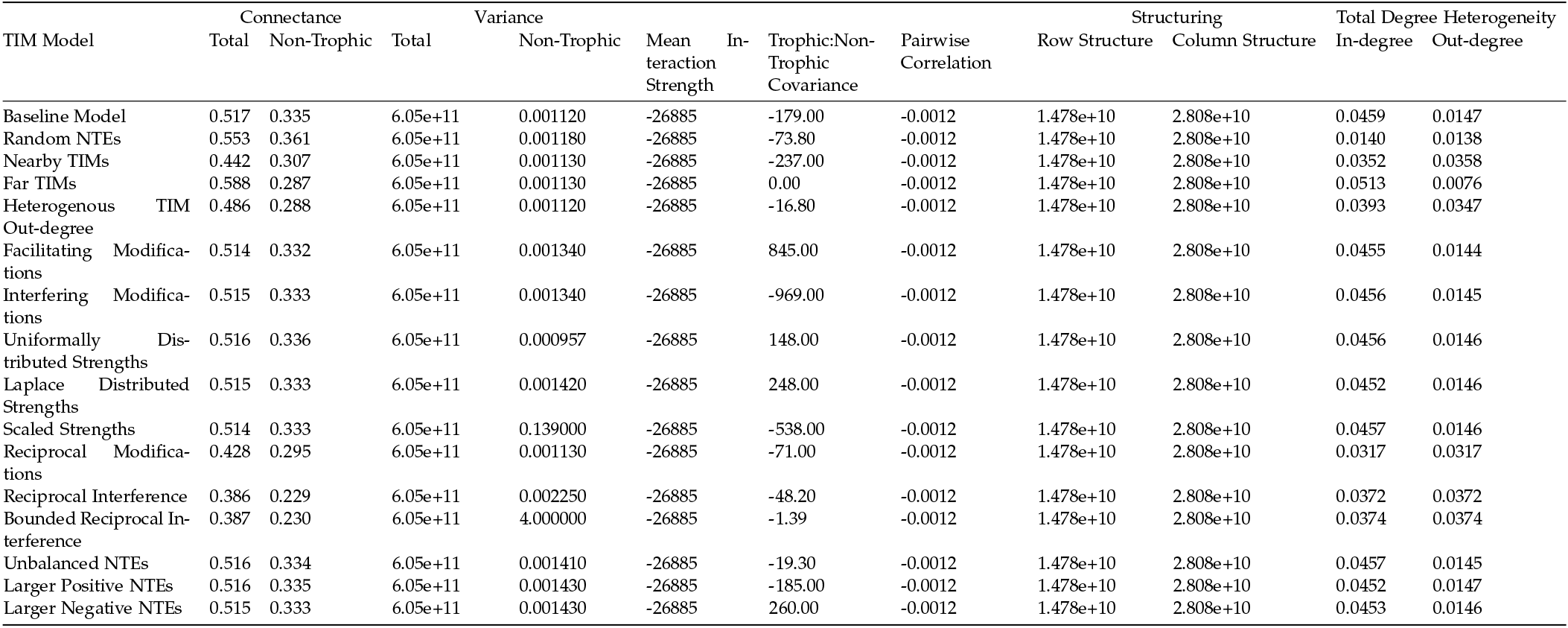
Table showing structural properties of Tampa Bay network with added TIMs.

## 6: Non-highlighted TIM Models

**Figure S5:**
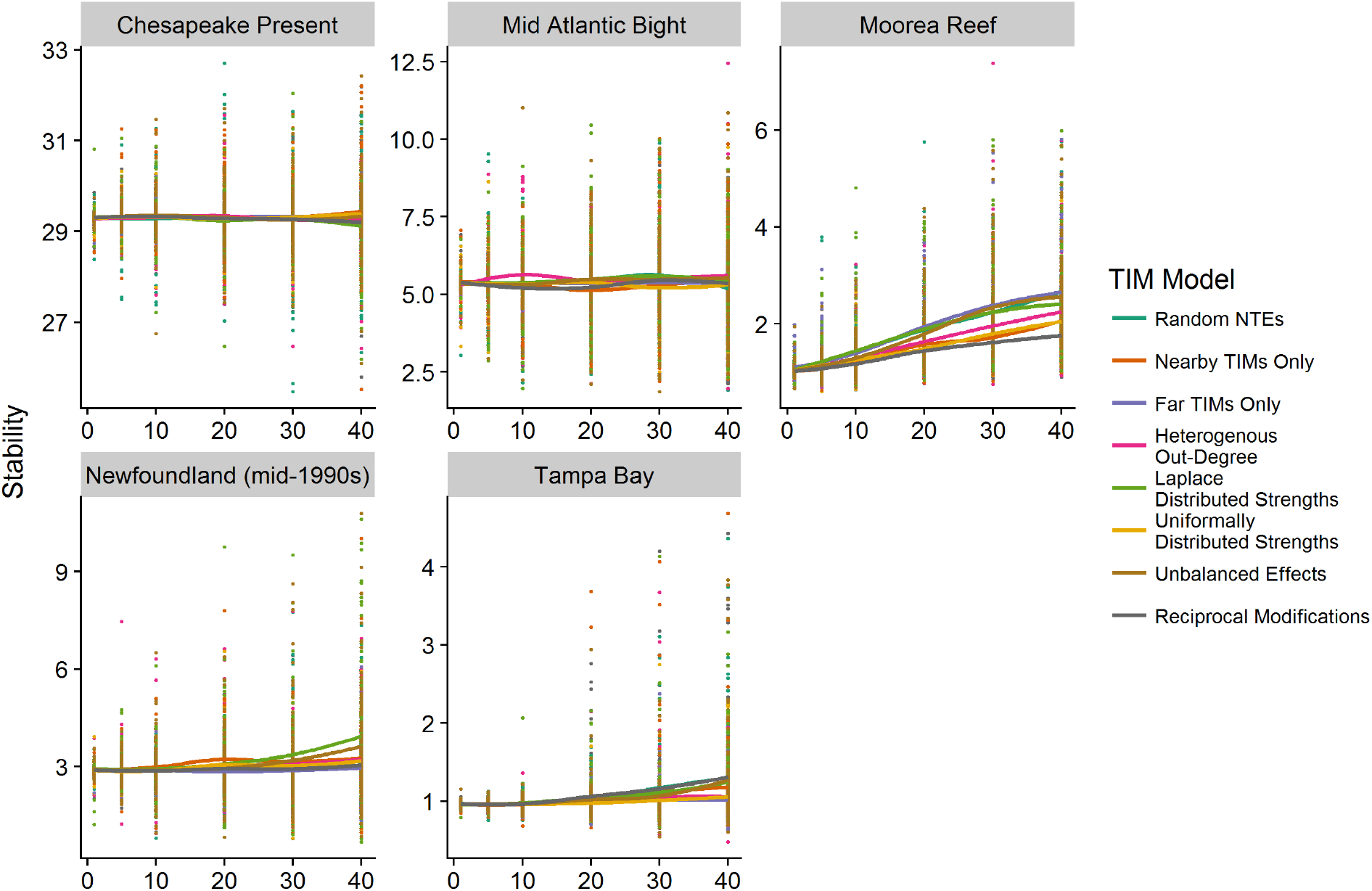
Stability response of the introduction of TIMs in the empricial webs under the models not highlighted in main text Figure 5.

## 7: Stability Criteria in the Empirical Networks

**Figure S6:**
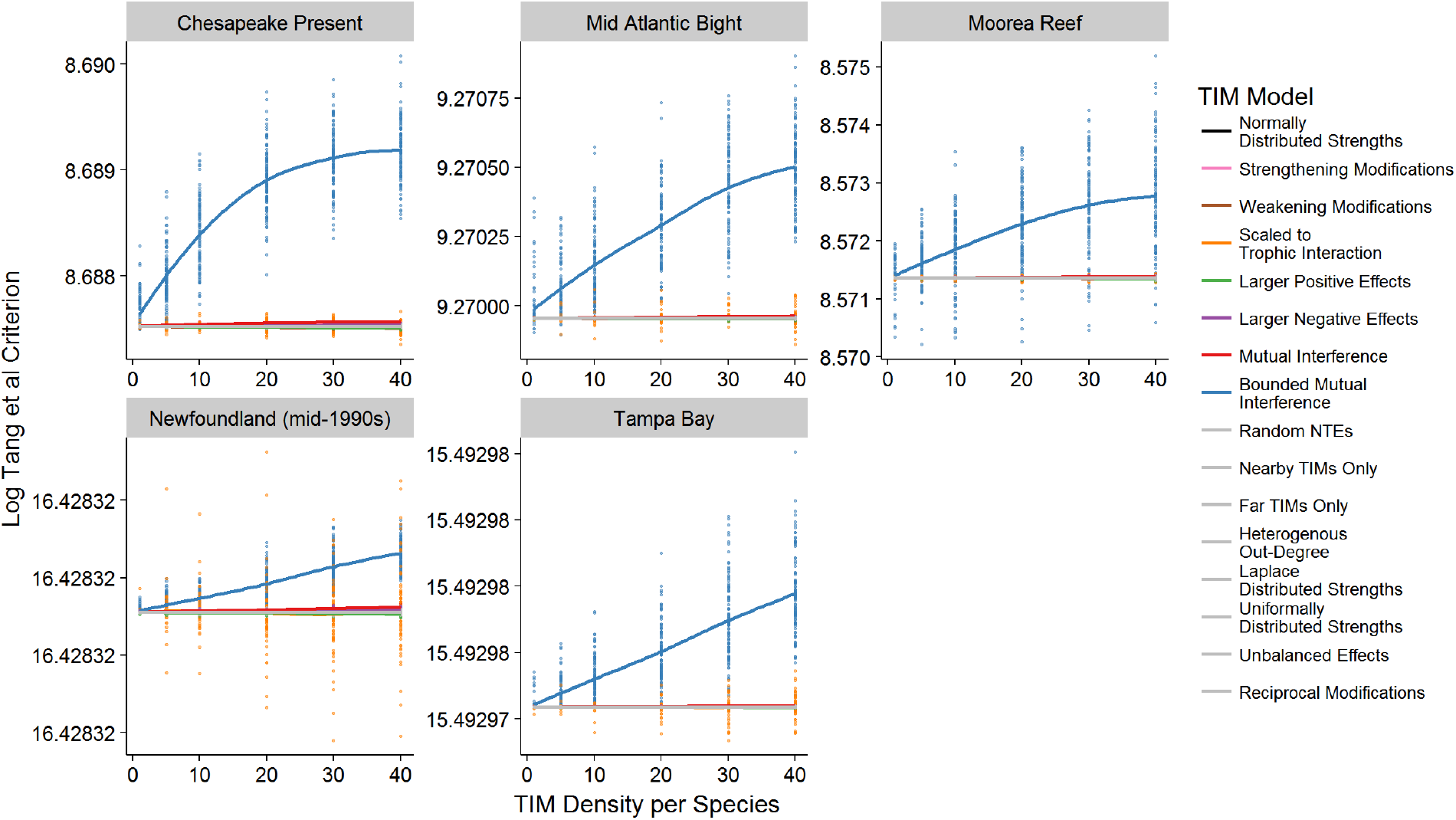
The minimal response of the Tang *et al.* stability criterion to the introduction of TIMs. Note the very restricted y-axis of each facet.

## Bibliography

Abrams, P. A. 1983. Arguments in Favor of Higher Order Interactions. The American Naturalist 121:887–891.

Abrams, P. A.. 2010. Quantitative descriptions of resource choice in ecological models. Population Ecology 52:47–58.

Allesina, S., J. Grilli, G. Barabás, S. Tang, J. Aljadeff, and A. Maritan. 2015. Predicting the stability of large structured food webs. Nature Communications 6:7842.

Allesina, S., and S. Tang. 2012. Stability criteria for complex ecosystems. Nature 483:205–208.

Allesina, S., and S. Tang. 2015. The stability–complexity relationship at age 40: a random matrix perspective. Population Ecology 57:63–75.

Arias-González, J. E., B. Delesalle, B. Salvat, and R. Galzin. 1997. Trophic functioning of the Tiahura reef sector, Moorea Island, French Polynesia. Coral Reefs 16:231–246.

Bairey, E., E. D. Kelsic, and R. Kishony. 2016. High-order species interactions shape ecosystem diversity. Nature Communications 7:12285.

Barabás, G., M. J. Michalska-Smith, and S. Allesina. 2017. Self-regulation and the stability of large ecological networks. Nature Ecology and Evolution 1:1870–1875.

Barbosa, P., J. Hines, I. Kaplan, H. Martinson, A. Szczepaniec, and Z. Szendrei. 2009. Associational Resistance and Associational Susceptibility: Having Right or Wrong Neighbors. Annual Review of Ecology, Evolution, and Systematics 40:1–20.

Brose, U. 2010. Body-mass constraints on foraging behaviour determine population and food-web dynamics. Functional Ecology 24:28–34.

Bruno, J. F., J. J. Stachowicz, and M. D. Bertness. 2003. Inclusion of facilitation into ecological theory. Trends in Ecology and Evolution 18:119–125.

Christensen, V., A. Beattie, C. Buchanan, H. Ma, S. J. D. Martell, R. J. Latour, D. Preikshot, et al. 2009. Fisheries ecosystem model of the Chesapeake Bay: Methodology, parameterization, and model exploration. NOAA Tech. Memo 1–146.

Christensen, V., and D. Pauly. 1992. ECOPATH II - a software for balancing steady-state ecosystem models and calculating network characteristics. Ecological Modelling 61:169–185.

Coyte, K. Z., J. Schluter, and K. R. Foster. 2015. The ecology of the microbiome: Networks, competition, and stability. Science 350:663–666.

de Ruiter, P. C., A.-M. Neutel, and J. C. Moore. 1995. Energetics, Patterns of Interaction Strengths, and Stability in Real Ecosystems. Science 269:1257–1260.

Delmas, E., U. Brose, D. Gravel, D. B. Stouffer, and T. Poisot. 2017. Simulations of biomass dynamics in community food webs. Methods in Ecology and Evolution 8:881–886.

Donohue, I., O. L. Petchey, S. Kéfi, A. Génin, A. L. Jackson, Q. Yang, and N. E. O’Connor. 2017. Loss of predator species, not intermediate consumers, triggers rapid and dramatic extinction cascades. Global Change Biology 23:2962–2972.

Dunne, J. A., and M. Pascual. 2006. Ecological Networks: Linking Structure to Dynamics in Food Webs. Oxford University Press, Oxford.

Fontaine, C., P. R. Guimarães, S. Kéfi, N. Loeuille, J. Memmott, W. H. van der Putten, F. J. F. van Veen, et al. 2011. The ecological and evolutionary implications of merging different types of networks. Ecology Letters 14:1170–1181.

García-Callejas, D., R. Molowny-Horas, and M. B. Araújo. 2018. Multiple interactions networks: towards more realistic descriptions of the web of life. Oikos 127:5–22.

Gibbs, T., J. Grilli, T. Rogers, and S. Allesina. 2017. The effect of population abundances on the stability of large random ecosystems. ArXiv 1708.08837.

Golubski, A. J., E. E. Westlund, J. Vandermeer, and M. Pascual. 2016. Ecological Networks over the Edge: Hypergraph Trait-Mediated Indirect Interaction (TMII) Structure. Trends in Ecology & Evolution 31:344–354.

Grilli, J., G. Barabás, M. J. Michalska-Smith, and S. Allesina. 2017. Higher-order interactions stabilize dynamics in competitive network models. Nature 548:210–213.

Grilli, J., T. Rogers, and S. Allesina. 2016. Modularity and stability in ecological communities. Nature Communications 7:12031.

Heymans, J. J., and T. J. Pitcher. 2002. A Model of the Marine Ecosystem of Newfoundland and Southern Labrador (2J3KLNO) In the Time Periods 1985-1987 and 1995-1997. Pages 5–43 in T. Pitcher, J. J. Heymans, and M. Vasconcellos, eds. Ecosystem models of Newfoundland for the time periods 1995, 1985, 1900 and 1450. (10(5).). Fisheries Centre Research Reports.

Ings, T. C., J. M. Montoya, J. Bascompte, N. Blüthgen, L. Brown, C. F. Dormann, F. Edwards, et al. 2009. Ecological networks - beyond food webs. Journal of Animal Ecology 78:253–269.

Jacquet, C., C. Moritz, L. Morissette, P. Legagneux, F. Massol, P. Archambault, and D. Gravel. 2016. No complexity–stability relationship in empirical ecosystems. Nature Communications 7:12573.

James, A., M. J. Plank, A. G. Rossberg, J. Beecham, M. Emmerson, and J. W. Pitchford. 2015. Constructing Random Matrices to Represent Real Ecosystems. The American Naturalist 185:680–692.

Jeschke, J. M., M. Kopp, and R. Tollrian. 2004. Consumer-food systems: why type I functional responses are exclusive to filter feeders. Biological Reviews 79:337–349.

Johnson, S., and N. S. Jones. 2017. Looplessness in networks is linked to trophic coherence. Proceedings of the National Academy of Sciences 201613786.

Jones, C. G., J. H. Lawton, and M. Shachak. 1994. Organisms as Ecosystem Engineers Organisms as ecosystem engineers. Oikos 69:373–386.

Kéfi, S., E. L. Berlow, E. A. Wieters, L. N. Joppa, S. A. Wood, U. Brose, and S. A. Navarrete. 2015. Network structure beyond food webs: mapping non-trophic and trophic interactions on Chilean rocky shores. Ecology 96:291–303.

Kéfi, S., E. L. Berlow, E. A. Wieters, S. A. Navarrete, O. L. Petchey, S. A. Wood, A. Boit, et al. 2012. More than a meal… integrating non-feeding interactions into food webs. Ecology Letters 15:291–300.

Kéfi, S., V. Miele, E. A. Wieters, S. A. Navarrete, and E. L. Berlow. 2016. How Structured Is the Entangled Bank? The Surprisingly Simple Organization of Multiplex Ecological Networks Leads to Increased Persistence and Resilience. PLOS Biology 14:e1002527.

Koen-Alonso, M. 2007. A Process-Oriented Approach to the Multispecies Functional Response. Pages 1–36 in N. Rooney, K. S. McCann, and D. L. G. Noakes, eds. From Energetics to Ecosystems: The Dynamics and Structure of Ecological Systems. Springer Netherlands.

Levine, J. M., J. Bascompte, P. B. Adler, and S. Allesina. 2017. Beyond pairwise mechanisms of species coexistence in complex communities. Nature 546:56–64.

May, R. M. 1972. Will a large complex system be stable? Nature 238:413–414.

May, R. M. 1973. Stability and Complexity in Model Ecosystems. Princeton University Press, Princeton.

May, R. M.. 1977. Predators that switch. Nature 269:103–104.

Mayfield, M. M., and D. B. Stouffer. 2017. Higher-order interactions capture unexplained complexity in diverse communities. Nature Ecology & Evolution 1:0062.

McCann, K., A. Hastings, and G. R. Huxel. 1998. Weak trophic interactions and the balance of nature. Nature 395:794–798.

Milo, R. 2002. Network Motifs: Simple Building Blocks of Complex Networks. Science 298:824–827.

Montoya, J. M., S. L. Pimm, and R. V Solé. 2006. Ecological networks and their fragility. Nature 442:259–64.

Mougi, A., and M. Kondoh. 2012. Diversity of Interaction Types and Ecological Community Stability. Science 337:349–351.

Murdoch, W. W., and A. Oaten. 1975. Predation and Population Stability. Pages 1–131 in A. MacFadyen, ed. Advances in Ecological Research Volume 9. Academic Press.

Neubert, M. G., and H. Caswell. 1997. Alternatives to resilience for measuring the responses of ecological systems to perturbations. Ecology 78:653–665.

Neutel, A.-M., J. A. P. Heesterbeek, and P. C. de Ruiter. 2002. Stability in Real Food Webs: Weak Links in Long Loops. Science 296:1120–1123.

Novak, M., J. D. Yeakel, A. E. Noble, D. F. Doak, M. Emmerson, J. A. Estes, U. Jacob, et al. 2016. Characterizing Species Interactions to Understand Press Perturbations: What Is the Community Matrix? Annual Review of Ecology, Evolution, and Systematics 47:409–432.

Ohgushi, T., O. Schmitz, and R. D. Holt. 2012. Trait-Mediated Indirect Interactions: Ecological and Evolutionary Perspectives. Cambridge University Press, Cambridge.

Okey, T., and R. Pugliese. 2001. A Preliminary ECOPATH Model Of The Atlantic Continental Shelf Adjacent To The Southeastern United Satates. Pages 167–181 in S. Guenette, V. Christensen, and D. Pauly, eds. Fisheries Impacts on North Atlantic Ecosystems: Models and Analyes (9th ed.). Fisheries Center Research Reports.

Olff, H., D. Alonso, M. P. Berg, B. K. Eriksson, M. Loreau, T. Piersma, and N. Rooney. 2009. Parallel ecological networks in ecosystems. Philosophical Transactions of the Royal Society of London B: Biological Sciences 364:1755–1779.

Pawar, S., A. I. Dell, and Van M. Savage. 2012. Dimensionality of consumer search space drives trophic interaction strengths. Nature 486:485–489.

Petchey, O. L., A. P. Beckerman, J. O. Riede, and P. H. Warren. 2008. Size, foraging, and food web structure. Proceedings of the National Academy of Sciences 105:4191–4196.

Pilosof, S., M. A. Porter, M. Pascual, and S. Kéfi. 2017. The multilayer nature of ecological networks. Nature Ecology & Evolution 1:0101.

Preisser, E. L., D. I. Bolnick, and M. F. Benard. 2005. Scared to death? The effects of intimidation and consumption in predator-prey interactions. Ecology 86:501–509.

Sander, E. L., J. T. Wootton, and S. Allesina. 2015. What Can Interaction Webs Tell Us About Species Roles? PLoS Computational Biology 11:e1004330.

Sanders, D., C. G. Jones, E. Thébault, T. J. Bouma, T. Van Der Heide, J. Van Belzen, and S. Barot. 2014. Integrating ecosystem engineering and food webs. Oikos 123:513–524.

Stouffer, D. B., J. Camacho, W. Jiang, and L. A. N. Amaral. 2007. Evidence for the existence of a robust pattern of prey selection in food webs. Proceedings of the Royal Society B: Biological Sciences 274:1931–1940.

Suraci, J. P., M. Clinchy, L. M. Dill, D. Roberts, and L. Y. Zanette. 2016. Fear of large carnivores causes a trophic cascade. Nature Communications 7:10698.

Tang, S., and S. Allesina. 2014. Reactivity and stability of large ecosystems. Frontiers in Ecology and Evolution 2:1–8.

Tang, S., S. Pawar, and S. Allesina. 2014. Correlation between interaction strengths drives stability in large ecological networks. Ecology Letters 17:1094–1100.

Tao, T. 2012. Topics in Random Matrix Theory. American Mathematical Society.

Terry, J. C. D., R. J. Morris, and M. B. Bonsall. 2017. Trophic interaction modifications: an empirical and theoretical framework. Ecology Letters 20:1219–1230.

Valdovinos, F. S., R. Ramos-Jiliberto, L. Garay-Narváez, P. Urbani, and J. A. Dunne. 2010. Consequences of adaptive behaviour for the structure and dynamics of food webs. Ecology Letters 13:1546–1559.

van Veen, F. J. F., P. D. van Holland, and H. C. J. Godfray. 2005. Stable Coexistence in Insect Communities Due to Density- and Trait-Mediated Indirect Effects. Ecology 86:3182–3189.

Walters, C. J., V. Christensen, S. J. Martell, and J. F. Kitchell. 2005. Possible ecosystem impacts of applying MSY policies from single-species assessment. ICES Journal of Marine Science 62:558–568.

Werner, E. E., and S. D. Peacor. 2003. A Review of Trait-Mediated Indirect Interactions in Ecological Communities. Ecology 84:1083–1100.

Williams, R. J., and N. D. Martinez. 2000. Simple rules yield complex food webs. Nature 404:180–183.

Williams, R. J., and N. D. Martinez. 2008. Success and its limits among structural models of complex food webs. Journal of Animal Ecology 77:512–519.

Wilson, D. S. 1992. Complex Interactions in Metacommunities, with Implications for Biodiversity and Higher Levels of Selection. Ecology 73:1984–2000.

Wootton, J. T. 1994. The Nature and Consequences of Indirect Effects in Ecological Communities. Annual Review of Ecology and Systematics 25:443–466.

Wootton, J. T., and M. Emmerson. 2005. Measurement of Interaction Strength in Nature. Annual Review of Ecology, Evolution, and Systematics 36:419–444.

Yodzis, P. 2000. Diffuse effects in food webs. Ecology 81:261–266.

